# FCHO controls AP2’s critical endocytic roles through a PtdIns4,5P_2_ membrane-dependent switch

**DOI:** 10.1101/2022.04.02.486817

**Authors:** Nathan R. Zaccai, Zuzana Kadlecova, Veronica Kane Dickson, Kseniya Korobchevskaya, Jan Kamenicky, Oleksiy Kovtun, Perunthottathu K. Umasankar, Antoni G. Wrobel, Jonathan G.G. Kaufman, Sally Gray, Kun Qu, Philip R. Evans, Marco Fritzsche, Filip Sroubek, Stefan Höning, John A.G. Briggs, Bernard T. Kelly, David J. Owen, Linton M. Traub

## Abstract

Clathrin-mediated endocytosis (CME) is the main mechanism by which mammalian cells control their cell surface proteome. Proper operation of the pivotal CME cargo-adaptor AP2 requires membrane-localised FCHO. Here, live-cell eTIRF-SIM shows that FCHO marks sites of clathrin-coated pit (CCP) initiation, which mature into uniform sized CCPs comprising a central patch of AP2 and clathrin corralled by an FCHO/Eps15 ring. We dissect the network of interactions between the FCHO interdomain-linker and AP2, which concentrates, orients, tethers and partially destabilizes closed AP2 at the plasma membrane. AP2’s subsequent membrane deposition drives its opening, which triggers FCHO displacement through steric competition with PtdIns4,5P_2_, clathrin, cargo and CME accessory factors. FCHO can now relocate toward a CCP’s outer edge to engage and activate further AP2s to drive CCP growth/maturation.

**125 character summary:** FCHO primes AP2 for CCV incorporation, a process that triggers FCHO release to enable activation/recruitment of further AP2s

## Main Text: Introduction

CME is the chief mechanism for swift and selective uptake of proteins into the intracellular endosomal system of eukaryotes (*1*). It is therefore key to controlling the plasma membrane proteome and thus cellular life. It is also the system that many invading pathogens, including most viruses, use for cellular entry and establishing infection (*2, 3*). The clathrin-coated vesicles (CCVs) that mediate CME, are formed from clathrin-coated pits (CCPs), which are scattered over the plasma membrane (PM) and can account for ∼2% of the cell membrane surface (*4, 5*). Clathrin, however, does not contact membranes and their embedded transmembrane protein cargo directly but is attached through membrane-bound clathrin adaptors. The principal endocytic clathrin adaptors are the heterotetrameric AP2 complex and monomeric CALM (*6*), both of which bind the defining marker of the PM, phosphatidylinositol 4,5-bisphosphate (PtdIns(4, 5)P_2_).

The AP2 complex, composed of *α*, *β*2, μ2 and *σ*2 subunits (Fig.1A), is pivotal to mammalian CCV formation (*7*) and its deletion is embryonically lethal in mice (*8*). Once recruited onto the cytosolic face of the PM, AP2 coordinates the assembly of a network of 300-400 proteins of ∼30 identities that comprise an endocytic CCP via an array of dynamic, µM K_D_ strength protein/protein interactions in a finely choreographed process that takes only 1-2 minutes (reviewed in (*5, 9–11*)). How, why and where CCPs are triggered to form on the PM are not clear. Cytosolic AP2 exists in a functionally closed, ‘locked’ conformation that is incapable of simultaneously utilizing its four PtdIns(4, 5)P_2_ binding sites to contact the membrane. The binding sites for both of its cognate Yxx*Φ* and [DE]xxxL[LI] cargo sorting signals are also occluded (*9, 12–15*) as is its primary LLNLD clathrin binding motif (*16*). To function in CME, AP2 must be both recruited to the PM and activated i.e. undergo a large-scale conformational reorientation of its subunits (*13, 16*) to permit full PtdIns(4, 5)P_2_, cargo and clathrin binding. Current models for CCV initiation propose the simultaneous arrival of the three pioneer proteins Eps15, FCHO and AP2 to form membrane-tethered nanoclusters (*17–19*) in which AP2 activation occurs via a molecular mechanism whose details are obscure.

**Fig. 1.**
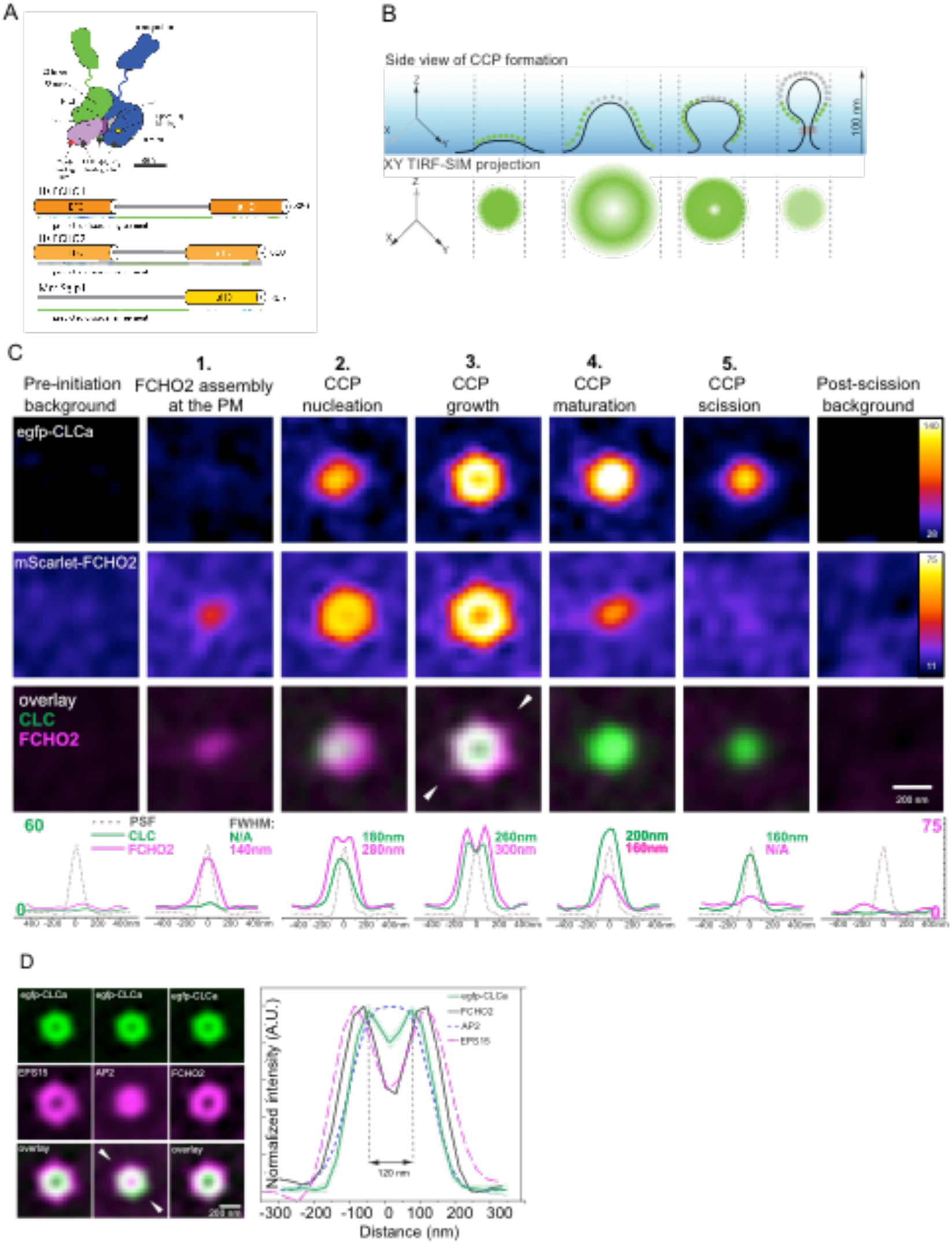
Dual colour live-cell microscopy with high-NA eTIRF-SIM reveals dynamic nanoscopic FCHO2 assembly that pre-determines site and size of clathrin coated pit formation. **A** Cartoon representations of mammalian AP2 and muniscin family members FCHO1, FCHO2 and SGIP. AP2 is coloured according to subunits as used throughout *α* (dark blue), *β*2 (dark green), Nµ2 (dark purple), Cµ2 (pale purple), *σ*2 (cyan). Muniscins possess FBAR and MHD domains joined by unstructured inter domain linkers **B** Upper panel: Side-view representation of a forming CCP imaged by eTIRF-SIM with an illumination depth of 100nm (blue gradient). Clathrin is depicted either in green, when visible and grey when not visible due to exponentially decaying evanescent wave of TIRF. Lower panel: Two-dimensional projection of clathrin (green) in a forming CCP along its axis of invagination into the plane of the plasma membrane. **C** eTIRF-SIM images of mScarlet-FCHO2^WT^ and egfp-CLCa in different phases of CCP formation. ROI containing single CCPs in a time series were dissected into phases depending on the appearance of FCHO2 or CLCegfp. ROIs with individual CCPs in different phases were assembled into a z-stack and averaged across 150 CCPs in 3 different cells. The bottom row shows comparison of not normalized intensity distribution profiles (along the line indicated with white triangles) for CLC and FCHO2 (green and magenta respectively) at different time points together with their full width at half maximum (FWHM). The black dotted line shows the measured point spread function (PSF). **D** Left Hand Panel: Averaged eTIRF-SIM images of multiple individual ROIs (as in C). The top row (green) shows CLC, the middle row (magenta) pioneer proteins of interest (Eps15, AP2 and FCHO2). The bottom row demonstrates overlay between the channels highlighting the molecule distribution with respect to CLC. mScarlet-FCHO2^WT^ forms annulus covering the outer rim of the clathrin hemispherical coat. of averaged CLC (green line) is 260nm. Right Hand panel: Normalized egfp-CLCa intensity profiles along the line indicated with white triangles of the previous panel. The green solid line shows the CCPs intensity profiles obtained from averaging images of for 480 CCPs in U-2 OS cells with estimated FWHM of 260nm. While AP2 (*α*-adaptin egfp, shown in blue dashed) forms a central patch in a mature CCP, Eps15 (poly clonal antibody, magenta dashed) forms a transition zone with FWHM larger than that of FCHO2 (black) and egfp-CLCa (green).

In vertebrates, there are three FCHO paralogues (muniscins): the *FCHO1* gene is expressed primarily in neuronal and lymphoid cells and *FCHO*2 is ubiquitously produced (*20–22*), while *SGIP1* expression is largely restricted to neuronal tissue (*23*). Complete *FCHO2* gene ablation in mice is post-partum lethal. Homozygous recessive *FCHO1* mutants cause immunosuppression and impaired clathrin-mediated endocytosis, which lead to death if not treated (*24, 25*). The reported effects on CME of FCHO depletion in cells have varied considerably from a relatively modest inhibition to complete block (*15, 18, 26-29*). FCHO1 and FCHO2 are similar in overall domain arrangement (Fig.1A) – folded N-terminal acidic phospholipid-binding homo-dimerization EFC/F-BAR domains (*30*) and C-terminal folded μ-homology domains (μHD) (*17*) separated by a ∼300 residue, low-complexity intrinsically disordered protein (IDP) linker, which has been suggested to participate in AP2 activation (*12, 15, 28*), although how it does so is undetermined. However, the importance of this linker to FCHO function was demonstrated by transfection of Cos7 cells depleted of FCHO2 by Sh RNA with a linker-deleted version of FCHO2 causing increased inhibition of TfR uptake as compared to the untransfected cells (*18*). Immuno- labelled deep etch-EM replicas (*17, 31*) first indicated FCHO/eps15 is localized mainly at the edges of flat, clathrin plaques, although controversies exist as to whether these are endocytically active (*32, 33*). Studies relying on analysis of fixed cells also reported the localisation of FCHO at the outer edge of CCPs in a ring (*18, 34*). However, due to the absence of a high-accuracy temporal dimension in combination with super-resolution spatial analysis, crucial questions remained unaddressed: These include does FCHO appear as a ring from the start of CCP formation? If not, then when does FCHO undergo annular reorganisation to the CCP periphery? Further, the distribution of AP2 in CCPs/CCVs remains unclear as it has been shown to be both present throughout them and at their edges (*34*) but also only on their membrane distal hemispherical parts (*35*).

Since it is of fundamental importance to understanding CME and thus to maintenance of the PM proteome, we set out to describe the spatial and temporal interrelationships between the key CME components AP2, FCHO/Eps15 and clathrin by using live-cell, high temporal accuracy, super-resolution imaging eTIRF SIM (enhanced Total Internal Reflection Fluoresence – Structured Illumination Microscopy). This achieved, we then deciphered the mechanisms of interaction with AP2 of four short, evolutionarily conserved blocks of sequence within FCHO linkers. X-ray crystallography, cryo-electron microscopy in solution (single particle analysis (SPA)) and on the membrane (tomography and subtomogram averaging (STA)) allowed us to understand how these interactions and their relation to AP2’s PtdIns4,5P_2_, cargo, clathrin and regulatory and accessory factor binding sites direct AP2’s PM deposition, activation and spatial separation from FCHO during CCP formation. Combining all of these data has allowed us to further develop the model for CCP initiation, growth and maturation at molecular resolution.

## Results

### Super-resolution dual colour live-cell microscopy reveals a critical role for dynamic FCHO2 assemblies in determining the fidelity of clathrin-coated pit formation

To understand how FCHO2 dynamic patterning drives CCP nucleation we used rapid live-cell multi-colour eTIRF-SIM and developed a semi-automated analysis pipeline of time-lapse movies (Fig. S1A), enabling us to resolve structures as small as 110nm simultaneously in all channels (Guo et al., 2018) (Figure 1B). We imaged the formation of CCPs in a U-2 OS cell line in which FCHO2 knockout was followed by stable reconstitution with near-endogenous levels of mScarlet-FCHO2WT (Figure S1 B,C,D) and stable expression of egfp clathrin light chain a (egfp-CLCa).

Based on the unique spatial and temporal patterns of FCHO2 in comparison to those of clathrin, we can identify five distinct CCP lifetime phases (Fig.1 B,C). In phase 1 we observed that FCHO2 forms a circular patch at the PM and its local concentration increases. During Phase 2 clathrin recruitment and then CCP nucleation begin. Automated quantitative analysis of conventional TIRF time-lapse movies revealed that 96% of all the dynamic and growing FCHO2 assemblies recruit AP2 and mature into CCPs (Fig.S2A,B,C). FCHO2 preceded AP2 and clathrin recruitment to the membrane as an FCHO2 signal was observed in the same pixel location at which a CCP finally appeared, prior to clathrin or AP2 signals. In agreement with Fig.1C, we observed that the FCHO2 signal reached its maximum intensity ∼10s earlier than the signals for AP2 and clathrin (Fig.S2B,C).

The semi-automatic quantitative analysis of super-resolved eTIRF-SIM movies (Movies S1, S2, S3, S4) revealed that during phase 2 where clathrin recruitment/CCP nucleation starts, the FCHO2 rapidly transformed from a patch into a ring as its abundance increases. To extract the spatial distributions of FCHO2 and CLCa, we plotted their relative radial intensity profile. The distance between the points where the intensity is half of the maximum is 35% larger for FCHO2 in comparison to CLCa. We interpret this as the FCHO2 patch redistributing to the periphery of the nucleating clathrin coated pit around the membrane invagination, which would be in line with fixed cell images from (*18, 34*). During Phase 3 (Fig. 1B,C) the FCHO2 ring expands further. The clathrin signal also appears annular, presumably due to the z projection of the spheroidal dome it must form (*36, 37*) and the exponentially decaying intensity of the incident light within evanescent TIR field.

We examined the spatial distribution of AP2, clathrin, FCHO and Eps15 in the CCP growth phase (phase3), identified on the basis of an annular reorganization of FCHO in live cells, by immunofluorescence. We found AP2 limited to a central patch, presumably at the distal end of the CCP in line with (*35*). FCHO2 had been proposed to bind AP2 on the membrane to potentiate its cargo binding (*12, 15, 17*). However, Fig1D shows that the FCHO ring segregates from the central AP2 patch into a ring, with the FCHO intensity maxima ∼50nm peripheral to the AP2 signal maxima (Fig.1D). This leaves only a thin interface where direct interaction of FCHO2 with open, membrane attached AP2 would be potentially possible: on the inside of the FCHO ring. A second potential interface exists on the outside of the FCHO ring where FCHO can engage with incoming ‘cytosolic’ closed AP2s. These data suggest that the current proposed model for FCHO function is oversimplified. EPS15 forms a phase-separated ‘liquid’ patch with FCHO2 prior to CCP nucleation (*38*); however, its function and organization later in CCP lifetime is less well established. Fig1D shows that Eps15 co-localises with FCHO2 in the annulus around the central AP2 patch.

We observed that during maturation (Phase 4) the FCHO2 signal had already started diminishing and had completely dissipated prior to scission (phase 5), in line with Merrifield’s early seminal work (*39*) and (*18*). However, since AP2 and FCHO2 have minimal, if any, interaction on the membrane later in a CCP’s lifetime as they have segregated into separate zones and there are only meagre quantities of FCHO2 but large amounts of AP2 in CCVs measured by quantitative mass spectrometry (*6*), they likely leave by different mechanisms: AP2 segregates to the distal end of the CCP, so dropping out of the evanescent field rather than leaving the CCP (*40*), whereas FCHO2 must diffuse from the CCP into the PM following competition for PtdIns4,5P_2_ by the arrival of late phase PtdIns4,5P_2_–binding endocytic proteins (*41*).

Finally, our TIRF data show an inverse correlation between the amount of the early FCHO2 accumulation and the time taken to eventual CCP scission (Figure S2A), suggesting the primary role for FCHO2 is in lowering the energetic and/or kinetic barriers for, i.e facilitating, CCP formation, presumably through its interaction with AP2 (*12, 15, 17*). The data also show that there is a striking size uniformity of all CCPs in both cell lines we tested: the average size of 480 circular CCPs did not deviate by more than 6% from an average diameter of 122 nm in U2-OS cells and RPE cells (Fig.1D). The calculated surface area is compatible with a resultant lipid vesicle diameter of ∼60 nm as is observed for CCVs isolated from cells (*11*).

Emerging from these observations is the crucial mechanistic question of how FCHO drives initial recruitment of AP2, yet subsequently segregates from AP2 (and clathrin) as the CCP grows when its role is meant to be activating AP2. Further, what interactions mediate the dynamic assembly and disassembly of the AP2-FCHO2-plasma membrane complex that permits the distinct rearrangements of these components throughout a CCP’s lifetime and how can FCHO facilitate CCP formation/AP2 activation?

### PtdIns4,5P_2_ membrane binding drives AP2 opening and competes off FCHO linker

The unstructured interdomain linkers of FCHO1, FCHO2 and Sgip1 have been proposed to interact with AP2 (*12, 15, 18, 42*). Bilayer interferometry (BLI) in near physiological conditions where AP2 core is known to be in its closed cytosolic conformation (*9*), showed the *K*_D_s for the direct interactions between AP2core and FCHO2 linker and between AP2core and FCHO1 linker were ∼10 μM (Fig.2A).

**Fig. 2.**
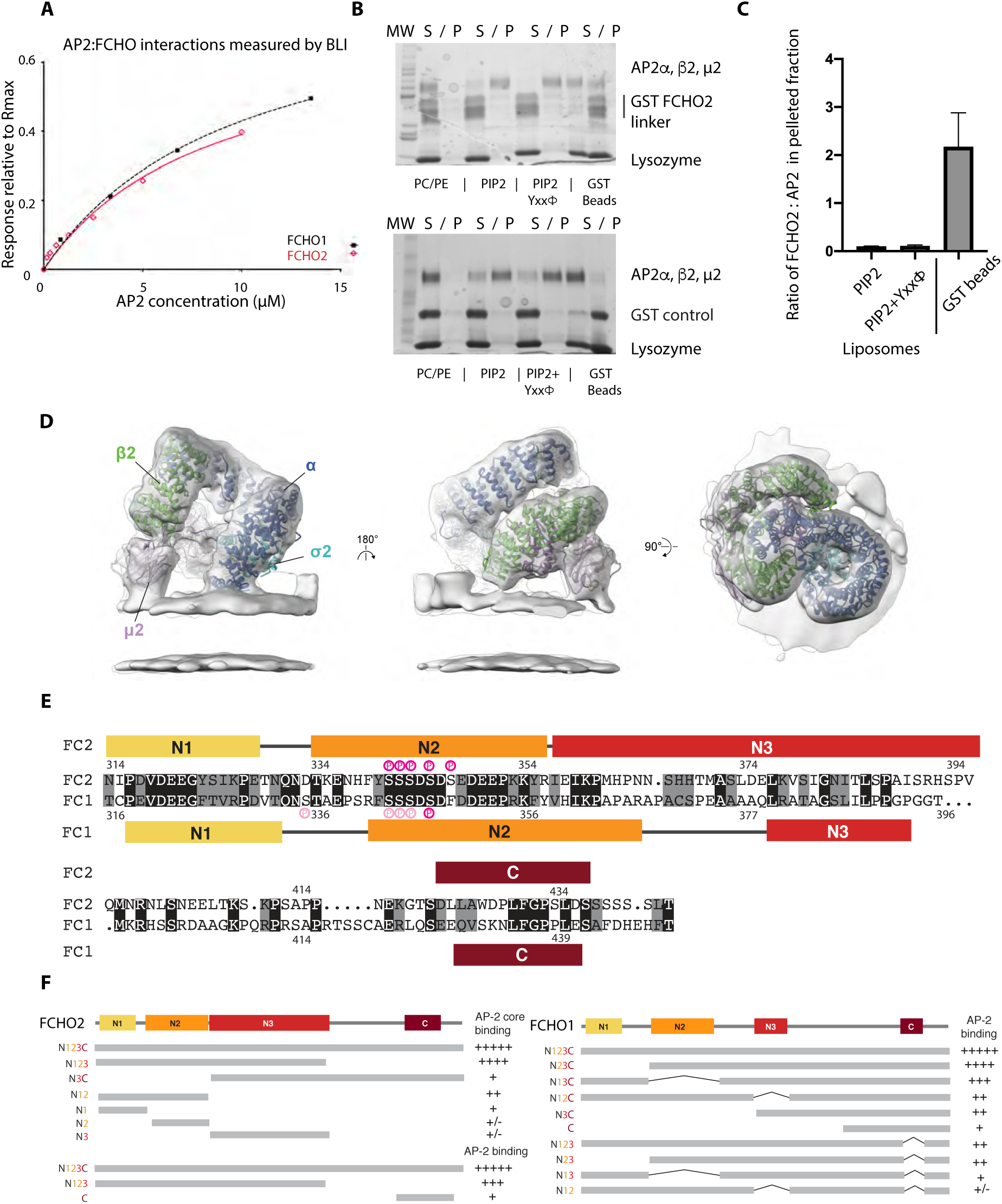
Four sequence blocks in the FCHO linker and PtdIns4,5P2-containing membranes compete for binding AP2. **A** Equilibrium analyses bio-layer interferometry (BLI) showing similar binding of recombinant AP2 core to FCHO1 linker (residues 316-444) K_D_8.5±2.2µM and to FCHO2 linker (residues 314-444) K_D_9.8±1.5µM. **B, C** Coomassie stained SDS PAGE gels of supernatant (S) and pelleted (P) fractions following addition of PC/PE, PC/PE/PtdIns(4, 5)P_2_, and PC/PE/PtdIns(4,5)P_2_+Yxx*Φ* cargo liposomes to mixtures of GSTFCHO2linker (15 µM) and AP2 core (1.25µM), which GSH sepharose bead pull downs show contain GSTFCHO2linker/AP2 core complex (top panel) and free GST and AP2 core (bottom panel). AP2 core binds efficiently to PC/PE/ PtdIns(4,5)P_2_, and PC/PE/ PtdIns(4,5)P_2_+Yxx*Φ*cargo liposomes, resulting in GSTFCHO2 present in preformed complexes being excluded to the soluble fraction since it contains no membrane binding function. **C** Quantitation of three independent experiments in **B** showing that the ratio of FCHO2:AP2 is ∼20 fold higher on beads than on PIP2 containing liposomes. **D** Tomographic structure of AP2 on the membrane in the presence of the FCHO2 linker. The cryo-EM map of AP2 recruited to cargo-free membranes in the presence of the FCHO2 linker, resolved to 9.7 Å, is shown as a grey isosurface. The map is fitted with the previously published ribbon model of AP2 bound to YxxΦ motif containing membranes in the absence of the FCHO2 linker (PDB: 6YAF), Here and throughout subunits are coloured *α* (dark blue) *β*2 (green) Nμ2 (dark purple) Cμ2 (pale purple) *σ*2 (cyan). **E** Human FCHO2 (FC2 upper) and FCHO1 (FC1 lower) inter-domain linker sequences aligned with identities highlighted in black and conservation in grey. Consensus CK2 phosphorylation sites are shown in dark pink and hierarchical CK2 sites in light pink. The conserved sequences blocks N1, N2, N3 and C assessed on the basis of conservations between sequences from across species and in the case of FCHO2 also on the basis of structure are coloured yellow, orange, red and claret respectively (colouring maintained throughout all subsequent figures) **F** Summary of the binding of the deletion constructs indicated of the unstructured FCHO2 and FCHO1 linkers to recombinant AP2 core and AP2 from brain cytosol as indicated. Gels shown in Figure S3

Next, we investigated AP2 membrane recruitment in the presence of the FCHO linker. Addition of PtdIns(4, 5)P_2_ and PtdSer or PtdIns(4, 5)P_2_ and PtdSer+YXXØ cargo liposomes to preformed AP2core•FCHO linker complexes in solution results in the AP2 binding to the membranes but causes the FCHO to become displaced from AP2 (in this assay, into the soluble fraction (Fig.2B,C)). These data indicate that FCHO linkers do not bind to membrane-associated AP2; that is, membrane and FCHO compete for binding to AP2. Nevertheless, FCHO2 linker does cause small but reproducible and significant increases in the steady state efficiency of AP2 membrane recruitment (∼13% for PtdIns(4, 5)P_2_ only liposomes and ∼22% for liposomes containing both PtdIns(4, 5)P_2_ and cargo) (Fig.S3A).

Cryo-electron tomography (Fig.2D) illustrates that when AP2 is added to liposomes containing PtdIns(4, 5)P_2_ and PtdSer but devoid of cargo, AP2 still assumes an open conformation, with PtdIns(4, 5)P_2_-binding BR3 and BR4 basic patches on Cμ2 (*43*) and the N-termini of *α* and *β*2, all simultaneously bound to the membrane surface (Fig. S3B,C,D). This conformation of AP2 is open and identical to the structure it adopts when cargo is also present in the membrane (*9*). It demonstrates that interaction with a high concentrations of free PtdIns(4, 5)P_2_/PtdSer alone is sufficient to drive AP2 into an active conformation. When a five-fold molar excess of FCHO linker is also included in the preparation of a cryo-ET sample, the same open conformation of AP2 is obtained, but even at subnanometer resolution there appears to be no FCHO linker electron density, in line with liposome pull downs (Fig.S3B,C,D). These data are supported by the absence of FCHO in mature CCVs (*6*) and our observed segregation of AP2 and FCHO during CCP maturation.

Taken together, these data indicated that FCHO2 binds AP2 in solution (in its closed, inactive conformation) to potentiate AP2 membrane deposition, but the two do not remain bound to each other on the membrane.

### Identifying and dissecting the AP2 binding sequence blocks in FCHO linkers

We set out to understand the molecular mechanism of FCHO•AP2 interaction and to explain why FCHO apparently does not bind AP2 on the membrane and hence we see them largely segregate in cells during CCP formation. The ubiquitously expressed FCHO2 (*21*) was used as it is less prone to degradation.

Phylogenetic analyses of FCHO linkers identified four blocks of potential functional importance on the basis of conservation of sequence and secondary structure prediction: in FCHO2, these were designated as N1 (314–328), N2 (334–357), N3 (358–397) and C (423–438) (Fig.2E,S3E). Single blocks or pairs of blocks showed little detectable binding to AP2 cores in near physiological conditions (Fig.2F,S3F) and removal of C block has little inhibitory effect. Binding of FCHO2 linker to full AP2 from brain cytosol was affected by C block deletion whilst isolated C block alone showed some binding, which hinted at the C block being involved in interacting with AP2 appendages (see later). Broadly similar results were obtained with FCHO1 (Fig.2F, S3G,H). Moreover, a synergistic (more than additive) association of AP2 with GST-FCHO1 N1+N2+N3 and GST-FCHO1 C, or between GST-FCHO1 (N2+N3) and GST-FCHO1 C occurs when both blocks are co-immobilized together upon GSH-Sepharose (Fig.2F, S3I).

### Closed AP2 binds FCHO2 linker

We were however unable to crystallize AP2 core in the presence of free FCHO2 linker. To increase the effective K_D_ of the interaction between FCHO2 linker and AP2 shown by BLI to have a K_D_ of ∼10µM and thereby improve the chances of isolating crystals of the complex, a number of chimaeras were made consisting of different combinations of blocks attached to different AP2 subunits. Most resulted in protein with poor purification properties and failed to provide material suitable for crystallization. However, one, a chimaera of AP2 core with FCHO2 N3-C region fused to the *β*2 C-terminus (AP2:*β*2FCHO2-N3+C) was crystallized in the presence of the PtdIns4,5P_2_ analogue D-myo-inositol-1,2,3,4,5,6-hexakisphosphate (InsP6), diffracted to 2.9-Å resolution and was solved by Molecular Replacement (MR) (*44*) (Fig.3A, S4A,B). AP2 was in its closed, inactive, conformation (not unexpected as no YXXØ cargo peptide was present). A predicted helix from the N3 block occupied the hydrophobic groove formed between *α*-helices 26 and 27 of *β*2-trunk (Fig.3A, S4B). The interaction is mediated by hydrophobic side chains from FCHO2, the pattern of which as well as a helical prediction (*45*) are conserved in FCHO1 and in Sgip1 N3 sequences (Fig.2E, S3E). Extrapolating from the finding that N3 must provide a major contribution to any FCHO•AP2, we studied the effect of deleting N3 in vivo on CCP phenotype *in vivo*. In line with the structure, whilst a construct comprising the FBAR and linker of FCHO can rescue the abnormal CCP phenotype seen in 1E HeLa cells (*15*) similarly to wt full length GFPFCHO, a GFPBAR+linker construct in which N3 is replaced with a similarly sized linker rich in glycine, serine and alanine residues cannot rescue the phenotype (Fig.S4C).

**Fig. 3.**
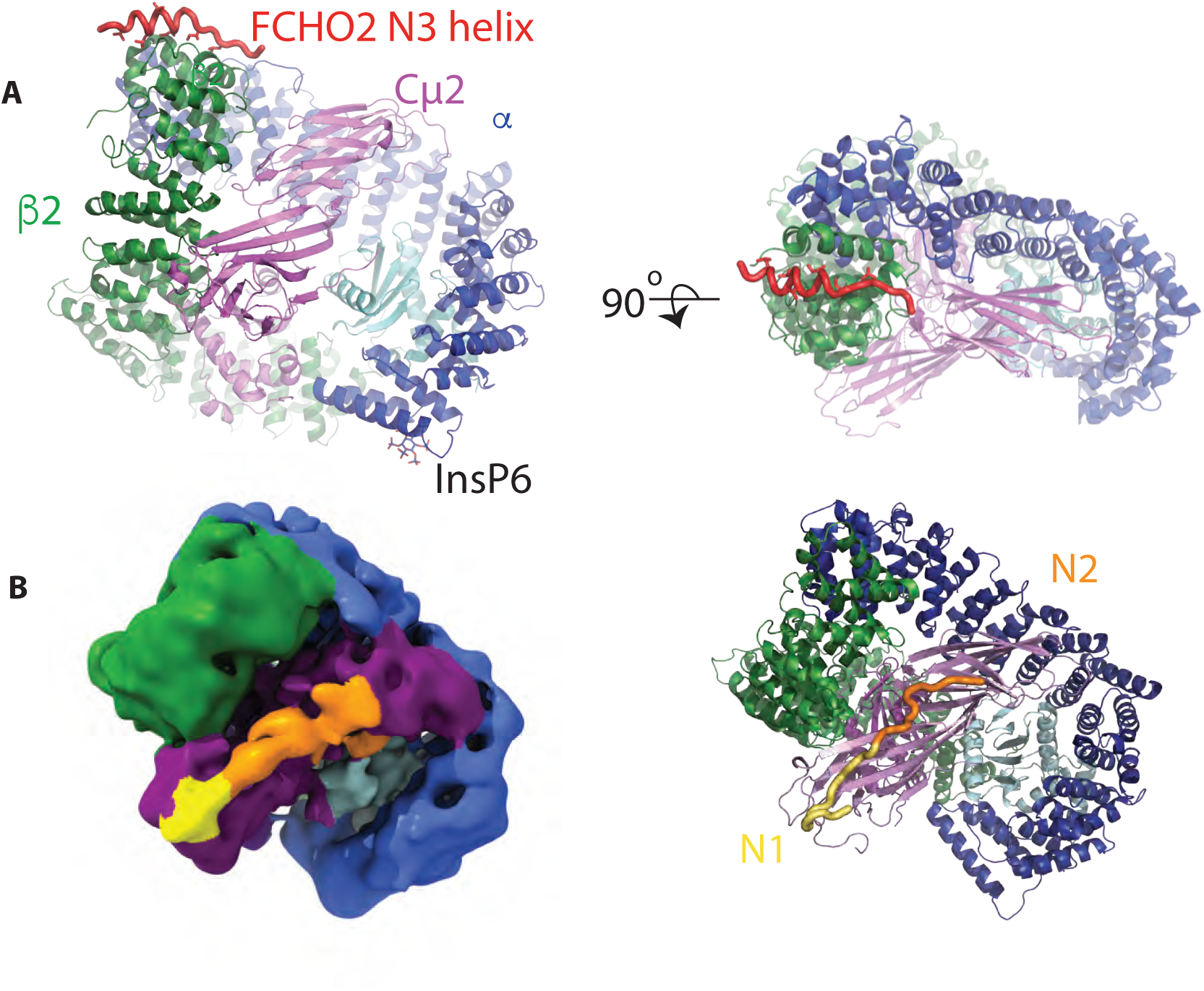
FCHO linker N1, N2 and N3 blocks can bind and activate closed and part open conformers of AP2. **A** Structure of AP2:*β*2FCHO2linker chimaera showing the position of the bound N3 helix (red) and InsP6 in orthogonal as indicated. **B** Left: Single particle EM reconstruction of AP2:µ2FCHO2-N1+N2+N3 chimera highlighting the position of FCHO2 N1+N2 (see N1N2-enriched subclass in Fig. S5 and Table S6). The map, locally filtered with a global resolution of 7 Å, is coloured according to AP2 subunit, the positions of which are derived from a fitted closed AP2 core (PDB ID 2vgl) and the approximate locations of N1 (yellow) and N2 (orange) are also indicated. N1 is persistent in all reconstructions of AP2:µ2FCHO2-N1+N2+N3 chimera, whereas N2 appears to be less fixed (Fig. S5): N3 is not seen in any single particle reconstructions. The position of N1 is in agreement with X-ray crystallographic structures (Fig. 4). N2 appears to extend from the N1 density associated with the Cµ2 BR3 site across the broad, positively charged BR4 surface patch of Cµ2. Right: Ribbon representation of AP2 in same orientation as left panel with FCHO linker shown in worm representation.

We turned to cryo-electron microscopy (single particle analysis, SPA) to try to resolve further details of the interaction. A chimaera in which N1-N3 of the FCHO2 linker was fused to the C-terminus of Cµ2 via a 30 residue linker (AP2:µ2FCHO2-N1+N2+N3) was suitable, and was used for SPA structure determination. A His_10_ affinity tag placed at the C-terminus of N3 allowed us to ensure that purified material contained full length N1-N3 linker using a NiNTA agarose recapture step. The resulting chimaeric µ2+N1 N2 N3 band was of the correct molecular weight and showed no degradation (Fig.S4D). 83% of the refined AP2 complex particles were used to determine a majority class structure at a resolution of 4 Å (Table S6): its conformation was closed (Fig.S4E,F,G) closely resembling the previously reported closed structure of AP2 core in physiological buffer (EMDB 10747, PDB 6yae). Reconstructions of several conformational subclasses were made (Fig. S5): the subclass with the clearest FCHO linker EM density, which was determined using ∼1/3rd of the particles used in the 7 Å structure determination, is shown in Fig. 3B. Extra features were clearly present in the EM map near the BR3 PtdIns4,5P_2_ binding site on Cμ2 (*43*), as well as also being observed near the BR4 PtdIns4,5P_2_ binding site, which we surmised would likely be N1 and N2 respectively (Fig.3B, S4E,F,G). The lower resolution of the N1 and N2 density (Fig. S5) suggests some conformational flexibility and/or sub-stoichiometric binding. No EM density for N3 was present in any of our reconstructions

### Molecular mechanism of FCHO2 linker N-terminus binding to AP2 Cμ2

To confirm the identity of the extra, Cµ2-proximal electron density as the N-terminus of FCHO2 linker, chimaeras of Cμ2 alone with N1+N2 attached to the C-terminus via 30 residue unstructured synthetic glycine, serine, alanine-rich flexible linkers were then created, crystallised and their structures solved at resolutions varying from 2.0 Å to 2.6 Å by MR using an unliganded version of Cμ2 as a search model. Diffracting crystals of (Cμ2-FCHO2-N1+N2) with either a thrombin cleaved GST tag (Fig.4A, S5D,E,F) or an N-terminal His_6_ tag (Fig.S5A) were obtained in the absence of free YXXØ cargo peptide. The Y pocket is variously filled by parts of the tags of an adjacent Cμ2 molecule. Residues from N1 wound around the N-terminus of Cμ2 (Fig.4A,B, S5A,B). Muniscin-family conserved side chains make key interactions with Cμ2 residues 167-170 (Fig.2E,3B,S5B), which form the BR3 PtdIns4,5P_2_ binding site (*13, 43*). ITC measurements (Fig.4C) showed the binding of FCHO2N1+N2 to have a K_D_ of ∼25 μM. Mutation of key binding residues in Cμ2 (E320A+E321A+K326A) reduced binding 5-fold and mutating Tyr323 to serine rendered it unmeasurably weak. The FCHO linker does not directly interact with the Y or Ø cargo motif binding pockets in any structure, but does pass close by before its main chain changes direction at FCHO2 Pro327. The weak density that follows on from N1 adopts a range of conformations forming crystal packing contacts, which we believe to be non-physiological (mutations FCHO2 Y340S and E336A+F339S have little effect on N1-N2 binding).

**Fig. 4.**
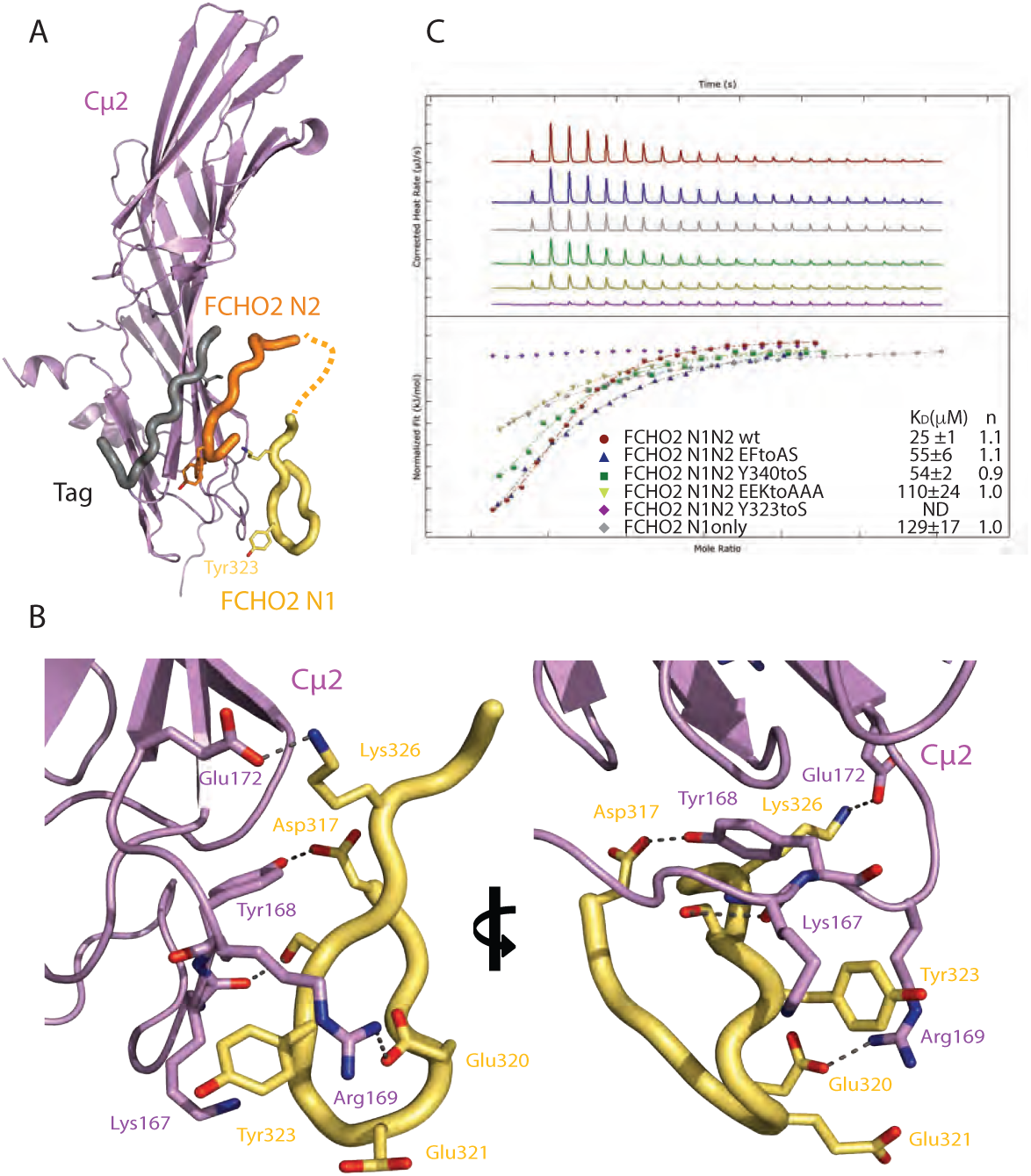
Binding of N1 to Cµ2. Overall positioning (**A**) and molecular details (**B**) of the N1 block bound to isolated Cμ2, allowing better definition of the N1’s binding to Cμ2 in whole AP2. The position of N2 and an affinity tag (grey) occupying the Yxx*Φ* binding site are also shown. The N1 interaction occurs mainly via complementary charged interactions between Asp318, Glu320 and Glu321 and the Cµ2 BR3 PtdIns4,5P_2_ binding site containing Lys167, Tyr168 and Arg169. **C** Confirmation of N1 binding site on Cμ2 by ITC using structure directed mutants. E321A+E322A+K326A reduces binding ∼5 fold from ∼25 μM and mutating Y323S renders binding unmeasurably weak and not determinable (ND). Mutations that fall outside the binding interface Y340S and E336A+F339S have little effect in binding. Deleting N2 also reduces binding 5 fold.

### Binding mechanism of poly acidic N2 to BR4 PtdIns4.5P_2_-binding site on µ2

Whereas N1+N2 binds Cµ2 with a K_D_ of around 25µM, N1 alone (i.e. deletion of N2) reduces binding by ∼5 fold (Fig.4C) demonstrating that N2 plays a role in binding AP2. The short, acidic N2 segment contains six glutamate/aspartate residues interspersed with serines, which are phosphorylated CK2 sites (*46*) (Fig.2E). These serine residues were mutated to glutamate to create a phosphomimetic version of the FCHO linker. (Fig.5A,B) show that both GST·FCHO2 wt linker and its phosphomimetic version are able to bind to AP2 and to isolated Cµ2, with the glutamate substitution markedly increasing binding, suggesting that N2 binds to a basic patch. Since the EM density following N1 in the AP2:µ2FCHO2-N1+N2+N3 chimaera SPA (Fig.3B) is located adjacent to the highly positively charged region of Cµ2 that includes Lys341, Lys343, Lys345 and Lys354 (*47*) termed the BR4 PtdIns4,5P_2_ binding site (Fig.5C) (*43*), this density can be assigned as N2. In support of this, mutating four of BR4’s basic residues to alanines (BR4-) caused substantial reduction in binding of GST·FCHO2 linker to Cµ2 (Fig.5D): the lower level of binding reduction of Serinc3 loop results from binding Cµ2 only at BR4 and not to BR3 (*43*), whereas in FCHO2 N1 can still bind to BR3 even when the BR4:N2 interaction is disrupted. These data demonstrate that N1 and N2 can bind through largely charged interactions to the BR3 and BR4 PtdIns4,5P_2_ binding sites on Cµ2 when AP2 is in its closed conformation,

**Fig. 5.**
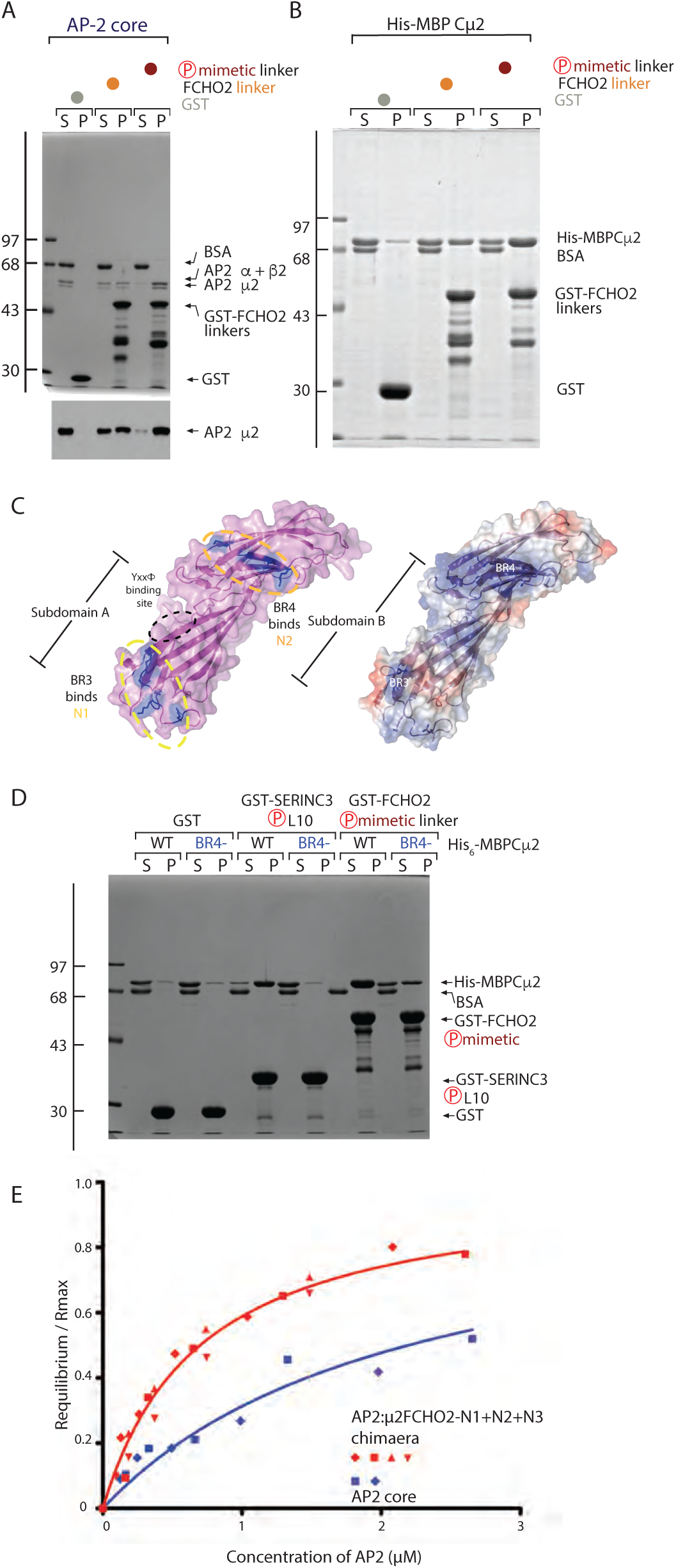
Binding of N2 to Cµ2 and AP2. **A** AP2 core binding to GST, GST-FCHO2 WT and phosphomimetic linker in pellet fraction (P). SDS-PAGE coomassie stained (top) and anti-μ2 subunit immunoblot (below). There is a clear loss of AP2core from the supernatant (S) fraction with either increased linker binding to phosphomimetic over WT FCHO2 linker in (P). **B** Affinity isolation of His-MBP-Cμ2 by GST, GST-FCHO2 WT and phosphomimetic linker in pellet fraction (P) shown by coomassie staining of SDS-PAGE. The extent of Cμ2 binding in the pellet (P) fractions is enhanced by S→E phosphomimetic substitution **C** Surface representations of the Cμ2 subdomain structure with positively charged residues from BR3 and BR4 highlighted in dark blue (left) and (right) same view of Cμ2 coloured for electrostatic potential contoured from −0.5 V [red] to +0.5 V [blue]. **D** WT or BR4- mutated (K339A,K341A,K343A,K345A) His-MBP-Cμ2 binding to GST, GST-SERINC3-L10 or GST-FCHO2 phosphomimetic linker as indicated by Coomassie blue staining of SDS PAGE. Robust binding of SERINC3-L10 or GST-FCHO2 phosphomimetic linker is shown in (P), which is severely inhibited by the BR4- compound mutation. Note the binding of GST-FCHO2 phosphomimetic linker is less affected than binding of SERINC3-L10 confirming the presence of another binding site for the FCHO2 linker on Cμ2 ie BR3 patch. **E** Binding of GST-Yxx*Φ* to AP2 core (blue) and AP2:µ2FCHO2-N1+N2+N3 chimera (red) in physiological buffer shown by BLI. AP2:*β*2FCHO2linker chimaera shows ∼3-fold tighter binding to TGN38 due to faster on- and slower off-rates than does AP2 core.

The binding of N1 to BR3 and N2 to BR4 combined with the fact that cryo EM tomography shows BR3 and BR4 sit directly on the membrane, provide the structural explanation for our observation that AP2 cannot simultaneously bind the FCHO2 linker and membrane-localised PtdIns4,5P_2_.

### Conformational destabilisation in AP2 is favoured by binding FCHO2

Compared to our previous analyses of AP2 by SPA EM in identical conditions (*9*), a much higher degree of particle heterogeneity was seen with AP2:µ2FCHO2-N1+N2+N3, likely indicating a decrease in stability of the closed conformation due to the presence of the FCHO linker. Previous crystallographic studies have shown that during the transition of AP2 between closed and open conformations, the µ2 subunit is repositioned and the gap between the *α*-solenoid stacks of the *α* and *β*2 trunks is reduced by ∼16 Å. In the range of particles observed of AP2:µ2FCHO2-N1+N2+N3, there is a contraction of the gap between the solenoids of ∼14 Å (Movie S5); yet without the movement of Cµ2 out of the AP2 bowl. A key residue known to latch β2 and Cu2 together in the phi pocket, *β*Val365, is no longer occluded in the FCHO-enriched SPA reconstruction of AP2:µ2FCHO2-N1+N2+N when compared to AP2 alone, further supporting a destabilisation of ’closed’ AP2 when FCHO2 is bound. Classification of the AP2:µ2FCHO2- N1+N2+N3 particles (see earlier) suggests that a significant minority of the particles adopt a ‘Cµ2- out’ conformation where the Cµ2 subunit is no longer contained within the AP2 bowl (Fig. S5) (that there was no degradation of µ2 was confirmed by SDS PAGE (Fig.S4D)). This conformation is reminiscent of that seen where Cµ2 is deleted and the AP2 bowl relaxes to the open conformation (*13*). Due to the high degree of flexibility we did not determine a higher resolution structure of this conformation. In conclusion significant conformational variability was present in the AP2:µ2FCHO2-N1+N2+N3 chimaera as compared to apo AP2 (*9*) with conformations ranging from closed through part open to a fully open bowl connected to a randomly positioned Cµ2.

We have been unable to visualize N1+N2+N3-simultaneously bound to closed AP2. This is despite firstly each block being able to bind individually to closed AP2 and secondly that in a model of such a complex (Fig.S6A) the distance FCHO needs to cross to link the Cμ2/N1 and *β*2/N3 binding sites intramolecularly (∼25 Å) is easily spanned by the intervening linker sequence. This could be explained by increased flexibility, conformational heterogeneity or both. A likely cause of this is that closed AP2 with simultaneously bound N1-N2-N3 is of higher energy than a number of more flexible, part open AP2 structures similarly bound to N1-N2-N3: such a range of structures would typically have Cµ2 at least partly displaced from the AP2 bowl but not into a fixed energy minimum position and hence we will now term them ‘not-closed’. Simultaneous binding of N1, N2, N3 may well also lower the energy barrier between closed and not-closed. The net result of both these processes would be to shift the equilibrium of AP2 core in solution away from closed. The energetic equilibrium between closed and open must be finely balanced as indicated by the distribution of particle conformers in SPA and be easy to ‘tip’ as witnessed by the increased protease sensitivity and rescue of FCHO deletion phenotypes in *C. elegans* (*12*) caused by introduction of any one of many mutations scattered all over AP2. This explanation is also borne out by our observation that all chimaeras containing AP2 core attached to a full length FCHO linker continuously lost material during purification unlike apo AP2 core, suggesting a destabilizing shift in equilibrium away from closed had occurred in these chimaeras.

In solution in 250mM NaCl, AP2 is closed and hence unable to bind N-terminal fluorescein- labelled Yxx*Φ* TGN38 peptide due to occlusion of its Cµ2 binding site by *β*2. Inhibition of binding can be relieved by the addition of ∼20-mer heparin to represent the highly negatively charged PtdIns(4, 5)P_2_ containing PM (Jackson et al 2010). Any conformation of AP2 in which Cµ2 is not tightly abutted against *β*2 trunk should be capable of binding Yxx*Φ* sequences. BLI (Fig.5E) indicates that in near physiological conditions, apo AP2 core must be somewhat destabilized as it shows some binding to GST-TGN38. In comparison, AP2:*β*2FCHO2linker chimaera showed ∼3- fold tighter binding to GST-TGN38 due to faster on- and slower off-rates: Note this is in line with speculations in (*12, 15, 17*) but is only relevant to a situation in solution, not *in vivo* where TGN38 cargo is membrane embedded. The shift in kinetics and thus binding caused by the presence of FCHO2 linker in solution is relatively modest but when combined with FCHO’s ability to concentrate AP2 at the surface of the plasma membrane, would bring AP2 membrane recruitment and cargo binding into the physiologically required time frame.

In considering the mechanism behind destabilisation of closed AP2 by the FCHO linker, the interface between Cµ2 and *β*2 is of particular interest since *β*2 residues Tyr405 and Val365 bind directly to Cµ2 at the positions occupied by the ‘xx’ and *Φ* residues respectively of cargo, and are key to holding AP2 closed (*14*). The position of FCHO N1 and N2 on Cμ2 and N3 on *β*2 trunk could function as provide anchor points on a closed AP2. This would position the linker between N1 and N2 such that it would clash with the backbone of the *β*2 400-410 loop, which is part of the *β*2:Cµ2 interface. Quite possibly that one ‘not-closed’ structure could actually be conformer we previously termed open+ (*48*). Using the experimentally derived positions of N1, and N3 *α*-helix binding sites on Cμ2 and *β*2, a model was created of open+ bound to the FCHO linker (Fig.S6B). In this, the distance between anchor points would again be ∼25Å and FCHO could be easily positioned to contact BR4 while the Yxx*Φ* cargo site would now be free and accessible. To reach the membrane-bound open conformation from ‘open+’, only a simple translation of 9 Å and a rotation of 90° of Cµ2 with respect to Nµ2 are required.

Thus, we propose that FCHO initially interacts with closed AP2, and in a mechanism likely driven by interference with the *β*2:Cµ2 interface, AP2 is shifted in its energetic landscape away from the closed conformation, so reducing the occlusion of the Yxx*Φ*-binding pocket on Cµ2 and facilitating its rapid opening by the planar PtdIns4,5P_2_-containing membrane: this agrees with our observation that FCHO potentiates the efficiency of AP2’s binding to membranes (see earlier).

### Structural analysis of FCHO2 linker binding to open AP2 in solution

Since FCHO destabilizes the closed conformation of AP2, TGN38 Yxx*Φ* cargo peptide (DYQRLN) was added in the hope of stabilizing an open form of AP2 in complex with FCHO2 linker. The resulting crystals diffracted to a resolution of ∼3.3 Å in space group P2_1_. The structure was solved by MR (*44*) using the bowl of the open AP2 and Cμ2 as sequential search models, and demonstrated that AP2 was in the open conformation (Fig.6A,B). In addition to clear electron density for Cμ2-bound YXXØ cargo and for the *σ*2-bound pseudo-dileucine cargo myc tag (Jackson et al., 2010), there was additional electron density for N3 assuming a helix of ∼3 turns occupying the same *β*2 trunk groove formed between *α*-helices 26 and 27 (Fig.6A,C, S6C) . Additionally, there was a short unstructured segment of peptide chain contiguous with it packing in the groove between helices 22 and 24. With the assistance of anomalous diffraction collected from crystals in which the FCHO2 linker was seleno-methionine-labelled, we were able to fully assign all of the N3 FCHO2 linker block so confirming our earlier identification of the N3 binding site on closed AP2.

**Fig. 6.**
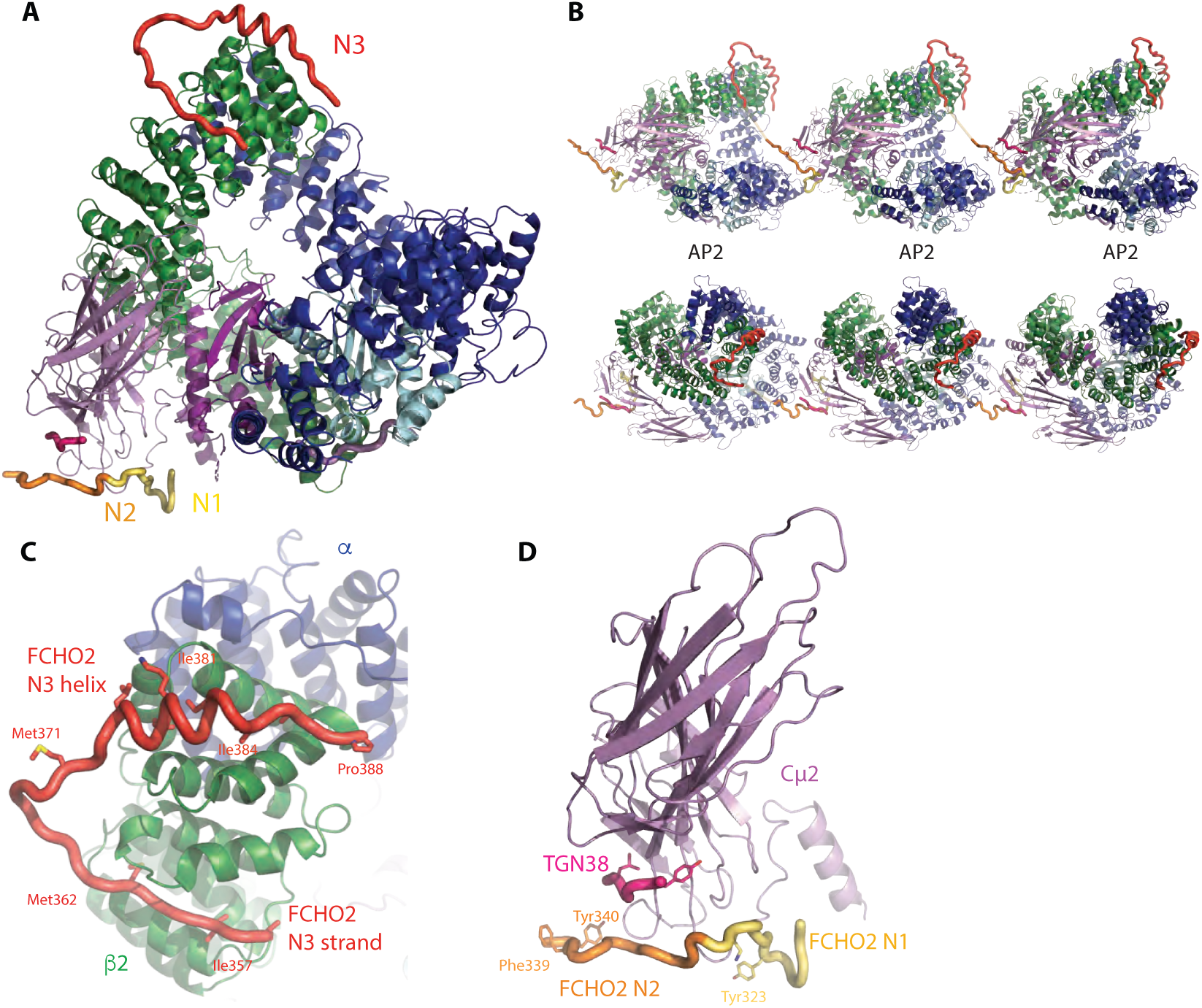
The binding of N1 and N3 FCHO blocks to open AP2. **A** Overall structure of a single AP2 core in its open form bound to N1 and N2 from one FCHO linker and N3 from a different FCHO linker (N1 yellow, N2 orange, N3 red, TGN38 cargo pink and in all subsequent images) **B** Orthogonal views of three AP2s in crystals formed in the presence of FCHO. The N1 and N3 blocks of any given linker bind to different AP2 molecules. N1 binds to the N-terminus of Cμ2 and N3 binds to the C-terminus of *β*2 from a different AP2. **C** Close up of the binding of N3 helix and preceding strand in hydrophobic troughs between helices 26 and 27 and between 22 and 24 respectively of *β*2 in open AP2/ FCHO2 linker complex crystals. **D** Close up of the position of the binding site of N1 and N2 to Cμ2 in open AP2/ FCHO2 linker complex crystals.

Transposing the position of N1 when bound to isolated Cμ2 onto this open AP2 core:FCHO2 linker complex structure demonstrates that the final piece of additonal electron denisty is N1 (Fig.6A,D, S6D) and that it binds in the same way in all the structures described in this work. The weaker electron density following N1, which ‘points’ at the start of an N3 attached to the *β*2-subunit of an adjacent AP2 molecule in the crystal, is the start of N2. Hence, the FCHO linkers appear to crosslink adjacent AP2s and thus provide a major crystal lattice packing interaction (Fig.6B). These data both confirm the locations of the N1 and N3 binding sites and suggest that if N1 and N3 are simultaneously bound as anchor points then the closed conformer becomes destabilised. However, since our earlier data also shows that binding of AP2 to membrane fully opens the AP2 and simultaneously outcompetes off the FCHO linker i.e. FCHO and hence membrane binding cannot be cotemporal, this solution structure cannot be physiologically relevant.Indeed, while the N1 and N3 binding sites are unobstructed in AP2’s open conformation, the distance (∼75 Å) that needs to be spanned is on the very edge of what is theoretically possible if all of the intervening sequence was completely extended - in practice this will not occur since even a glycine-serine peptide chains only exist in a state that is considerably less than "fully extended" (*49*). These observations, along with the absence of FCHO crosslinked and arrayed AP2s in cryo EM tomography and SPA, lends further suport to the conclusion that FCHO does not bind membrane-attached open AP2.

### Molecular mechanisms of FCHO2 linker’s C block binding to AP2’s *α*-appendage and Cμ2

The final highly conserved region of the FCHO linkers is block C, which contains an LFGPXL sequence (Fig.2E, S3E) separated from N3 by a variable-length (30-40 residue) disordered linker. Since FCHO1 and FCHO2 have been suggested to bind the AP2 appendages (*28*), we investigated this possibility.

ITC shows that various similar length control peptides containing several hydrophobic residues do not bind *α*-appendage, whereas the C block of FCHO1, binds with a *K*_D_ of ∼50 μM, whilst the FCHO2 and SGIP C blocks bind with a *K*_D_ of 100-200 μM (Fig.S7A). No binding was observed between the different C-blocks and the AP2 *β*2-appendage. Crystals of the *α*- appendage were grown in space group C 2 2 2_1_ in the presence of a 10-fold molar excess of FCHO1 C peptide (residues 426-448 SEEQVSKNLFGPPLESAFDHEH), which diffracted to 1.4Å resolution. The structure was solved by MR using an unliganded *α*-appendage structure (PDB 3hs8) as the search model. Density for the peptide was clearly visible and could be unambiguously built (Fig.7A, S7B). Although the peptide bound to the top platform site of *α*-appendage in the same position as FXDXF and DP[FW] motifs (*50–52*) the orientation is opposite to that in these previously determined structures (Fig.7B,C). The conserved hydrophobic interactions by the FCHO1 LFGPXL Phe436 and Leu440 (equivalent to FCHO2 residues 431 and 435) anchored the peptide to the *α*-appendage in the same positions as the phenylalanines of the FXDXF motif, facilitated in part by FCHO1 Pro438 being in a cis conformation: mutations in FCHO1 FGtoAA, LEStoAAA and PPtoGG reduce binding to *α*-appendage to below detectable levels (Fig.7D). H-bonds between the LFGPXL motif’s main chain and *α*-appendage side chains further stabilize the interaction. These observations suggest that muniscin LFGPxL ligands could be competed from the *α*-appendage during the later stages of CCP formation as the number and variety of accessory and regulatory proteins containing FxDxF and DP[FW] motifs increases (*39*)

**Fig. 7.**
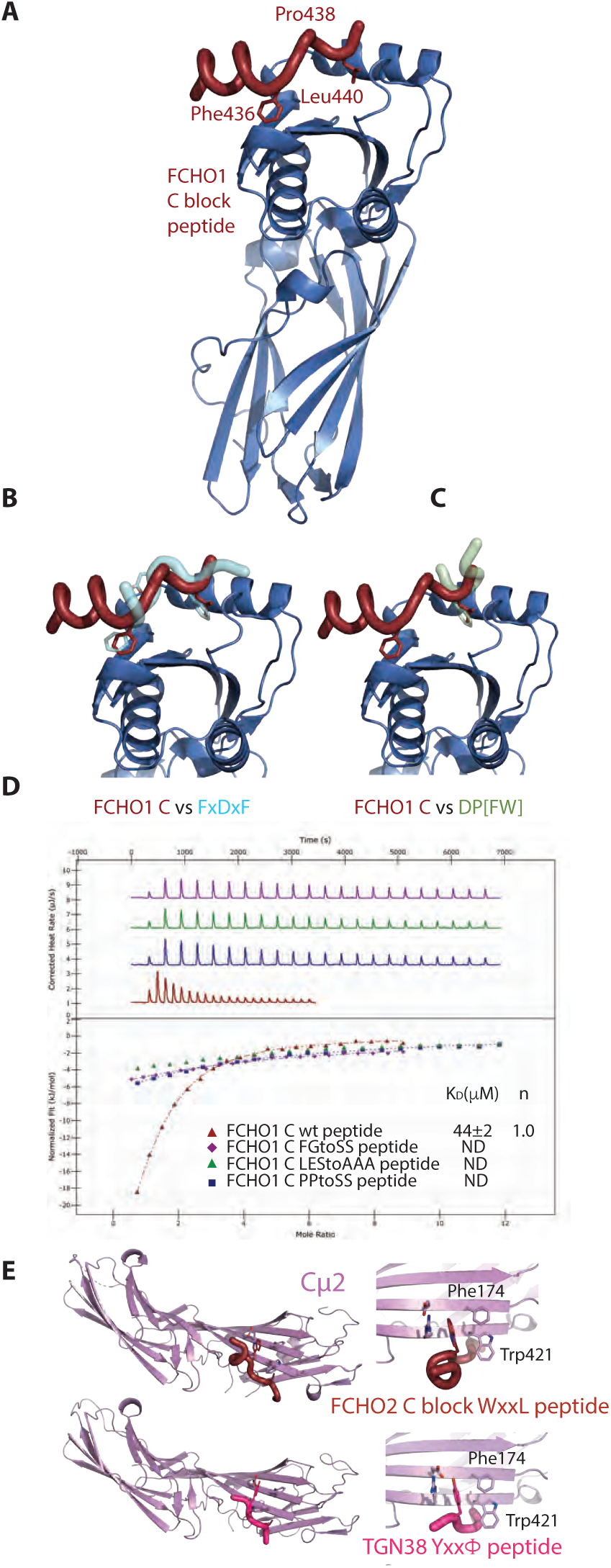
Alternative binding sites of the FCHO linker C block on α-appendage and Cμ2. **A** Structure of *α*-appendage (blue) showing the FCHO1 linker C block (LFGPPLES) bound to the platform subdomain at 1.4 Å resolution. Key binding residues are indicated. The residues preceding the core FGPPL motif for a short helix that packs against the side of the platform subdomain. **B, C** Superposition of FCHO1 linker C block with Left: bound FxDxF peptide (PDB 1KYU7) and right: bound DPF peptide (PDB 1KYU) both shown in semi-transparent representation. The FCHO1 SKNLFGPPLES peptide binds in the opposite orientation to FxDxF and DPF peptides whose final F and central F respectively sit in the same hydrophobic pocket as the C-terminal L of LFGPPL. **D** ITC showing ∼50 µM K_D_ binding of LFGPPL to *α*-appendage. Mutation of FG to SS, LES to AAA and PP to SS all reduce binding to weaker than 500µM K_D_. **E** Upper panels: 1.7 Å resolution Structure of FCHO2 WxxL motif of the WDPLFGPSLDS C block (claret) bound to Cµ2 at its Yxx*Φ* motif binding site (K_D_ 56µM and electron density see Figure S6) in perpendicular views. Lower panels same perpendicular views of Cµ2 bond to TGN38 Yxx*Φ* cargo peptide (pink)

Whilst FCHO2 clearly shows binding to *α*-appendage (Fig.S6A), we also noticed the presence of a highly conserved W preceding FCHO2’s LFGPxL by two residues. When combined with the first conserved L of the LFGPxL, this results in the creation of a WxxL sequence, which has been proposed as an alternative Cμ2-binding cargo sorting motif (*53*). An FCHO2 peptide incorporating this WxxL bound to Cμ2 in ITC with a K_D_ of ∼25μM (Fig.S6C), which is of similar strength of binding to many characterized Yxx*Φ* cargo motifs (*54*). Binding was dependent on the presence of the tryptophan side chain as the homologous sequence from FCHO1, which does not contain a tryptophan, showed no binding. Crystals of a complex of Cμ2 and a peptide corresponding to this WxxL signal were grown, diffracted to 1.7 Å resolution and were solved by MR using 1.9 Å unliganded Cμ2 as a search model. The backbone of the WxxL superimposes with that of a Yxx*Φ* peptide with the tryptophan side chain packing in the Y pocket and the leucine sitting in the *Φ* pocket (Fig.6E, S6D). These data suggest the possibility that the C block of FCHO2 has two possible modes of interaction with AP2: one with *α*-appendage and one with Cμ2. The mode favoured will likely be context dependent and it will likely be affected by multiple spatial and temporal factors in a nascent CCP including how many other *α*-appendage binding CCV proteins are present and whether the Yxx*Φ* site of Cμ2 is accessible and/or occupied by cargo. These binding modes are likely to be in dynamic equilibrium with each other. The absence of WxxL in the C block of neuronally-enriched FCHO1 but its presence in the ubiquitously expressed FCHO2 may reflect the paucity of Yxx*Φ* containing cargo found in synaptic CCVs (*55*) that could compete out WxxL from Cµ2, which if it did not occur could slow or stall CCV formation.

Thus, C blocks of all muniscin family members can bind to the *α*-appendage platform in a manner that is competitive with the binding of FxDxF and DP[FW] motifs of accessory CME proteins and in addition the partially overlapping WxxL sequence found only in FCHO2 can bind to the Cµ2 Yxx*Φ* cargo binding site, and therefore potentially stabilize the un-closed but not yet open form of AP2.

### The absence of the identified blocks in FCHO alters CCP formation dynamics

Taken together, these data suggest that the FCHO linker plays an important role in controlling AP2 during CCP formation. The AP2:FCHO interaction has a low µM apparent K_D_, but each of the four FCHO blocks’ individual interactions have only weak K_D_s i.e it works by avidity effects. This results in a highly dynamic, readily reversible, easily regulatable system. However, it also means that point mutation(s) in any one FCHO block will very likely only have minimal effects on the overall AP2:FCHO interaction. Indeed, complete deletion of an entire individual block has only minimal effects on FCHO’s AP2 binding *in vitro* (Fig. 2). Although replacing N3 with an equal length, unrelated sequence alters AP2 puncta morphology as in (*15, 17*) *in vivo* (Fig S4C), in order to see reliable effects on overall CCP behavior, and taking into account the redundant nature of interactions within the CCP component networks (*56*), we decided to investigate the effects of replacing the whole 140 residue FCHO linker with an equally unstructured, flexible, random sequence (FCHO2^STRING^) on CCP dynamics. Fig. S8A shows that transfection with this mutant, similar to the FCHO2 knockout in U2OS cells, leads to an increase in initiation rates of subthreshold ‘abortive’ clathrin positive structures i.e of less than 10s lifetime(*18*), when compared to rescue with wild type full-length FCHO2.

## Discussion

This work demonstrates how and why FCHOs underpin CCV formation. Nanoclusters of FCHOs linked by Eps15 (*17*) into phase separated micro-structures (*38*) bind, concentrate, orientate and activate AP2 at the PM, so potentiating AP2’s productive deposition on the PM to form the nucleus of a CCP. The multiple, dynamic interactions of short sequence blocks within the FCHO inter-domain linker with various AP2 subunits ensure that the process is of high fidelity and is sufficiently rapid, but is also readily reversible to allow proof-reading. Since association of AP2 with the membrane triggers the disassembly of AP2•FCHO complexes, this allows reuse of the same FCHO to activate further AP2s (i.e. FCHO could be considered effectively catalytic) to grow a CCP nucleus and in so doing provides both temporal processivity and spatial control to CCP formation. Thus, the making and subsequent breaking of the network of FCHO/AP2 interactions is key to the efficient, controlled genesis of endocytic CCVs and thus to the PM proteome. We can now describe an integrated mechanistic model for initiation of CCP/CCV formation *in vivo* (Fig.8) in which the actual mechanism of AP2 opening, driven by PtdIns4,5P_2_- binding and stabilized by cargo binding will be essentially the same *in vivo* and *in vitro* (described in (*13, 16*) but is facilitated by interaction with FCHO.

**Fig. 8.**
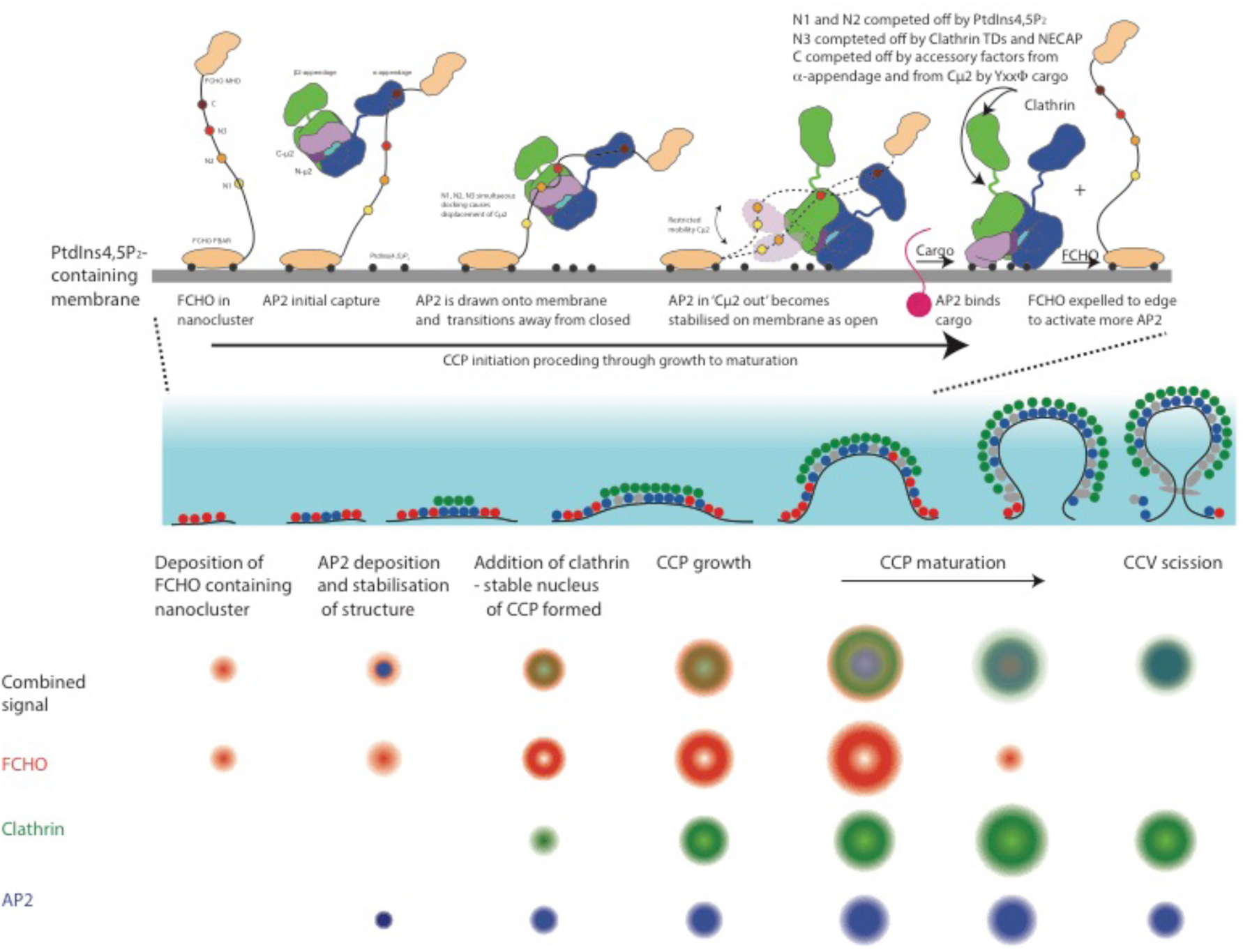
A molecular mechanistic model for the steps in the FCHO dependent activation of AP2 during CME. Top panel: FCHO2 bound to the membrane uses its four linker blocks, C, N3, N2 and N1 to engage with a closed, cytosolic AP2. Full binding results in the AP2 being orientated and concentrated on the membrane and destabilized to a ‘µ-out’ , flexible, conformation. This conformer quickly binds to the membrane causing it to fully open and N1 and N2 to dissociate. Cargo binding stabilizes this open form whilst the binding of polymerized clathrin and regulatory/accessory proteins competes out N3 and C and the FCHO is expelled to the edge of the forming CCP where it can recruit further AP2s. This process occurs during the phases of CCP growth indicated Middle panel: Schematic representation perpendicular to the PM of the phases of a CCPs life from initiation to scission: Left to right. FCHO2 red; AP2 blue; clathrin green; other clathrin adaptors and accessory protein grey. The pale blue gradient indicates an ∼100nm evanescent field as used for visualization in C Lower panel: Schematic representation of projections of a CCP lifecycle’s phases measured or inferred from eTIRF-SIM data. Top line – overlay of respective signals of FCHO2 (red); clathrin (green) and AP2 (blue).

### Interaction of AP2 with FCHO orients AP2 with respect to PM and promotes AP2 opening

Of the four FCHO linker blocks, N1 is nearest the PM as it follows directly after the FCHO FBAR domain with N2, N3 and C blocks increasingly distant from PM. As C can sample the largest volume to find AP2s, it seems likely that C block engagement will be the most frequent founder interaction. Although the order of engagement of subsequent linker blocks can be variable i.e is a fuzzy process, the most likely subsequent order would be N3, N2, N1 as this would correctly orientate, ‘draw down’ and hold an AP2 at the membrane surface, resulting in an AP2•FCHO complex structure resembling that modelled in Fig.S8B. Simultaneously, destabilization of AP2’s *β*2/μ2 interface (possibly enhanced by interactions of *α* and *β*2 with membrane PtdIns4,5P_2_), shifts the equilibrium of the AP2 away from fully closed. This increases the occupancy of ‘µ2 out’/‘not-closed’ conformations, in which the AP2 bowl has relaxed to a lower-energy, more open-like state (Fig. S5) and (*13*). Cµ2 can now sample its local environment on its flexible μ2 interdomain tether so that it can dock onto PtdIns4,5P_2_ via its BR3 and BR4 sites and become fully open. AP2’s subsequent local search dimensionality for cargo is effectively reduced from 3D to 2D: any cargo will be swiftly bound so further stabilizing an AP2’s open conformation and extending its PM dwell time. When stably open, AP2’s clathrin-binding *β*2-hinge is released (*16*) and so clathrin recruitment and polymerization can start: the process is enhanced by virtue of FCHO/Eps15 annular assemblages likely corralling open AP2, with which it will have little or no interaction, to increase its effective concentration and facilitate its crosslinking by a forming clathrin lattice. If sufficient AP2s become stably PM-attached through clathrin-crosslinking and cargo binding, the nucleus of a CCV has been formed.

### Interaction of AP2 with membrane and clathrin competes off FCHO interactions

Steric clashing causes competitive binding of PtdIns4,5P_2_ and FCHO N1/N2 blocks to Cµ2 BR3 and BR4 sites. This results in N1 and N2 being displaced as AP2 assumes a fully open conformation on the membrane and an FCHO•AP2 complex being significantly weakened as it can be mediated only by N3 interacting with *β*2-trunk and the C block with *α*-appendage or Cµ2. C binding will be subject to competition from incoming regulatory and accessory CCV proteins and other clathrin adaptors if it is bound to *α*-appendage; on the other hand, if it is bound to Cµ2, thus preventing AP2 reclosure and also Cµ2 binding to Yxx*Φ* motifs on cytosolic proteins, it will be expelled by Yxx*Φ* cargo binding. The N3 binding region on *β*2-trunk overlaps with the clathrin terminal domain layer in assembled clathrin coats ((*9*), Fig.S9A) which will likely serve to drive N3 off AP2. Intriguingly, a further source of competition may also exist for N3 binding to *β*2-trunk. A very similar sequence to the N3 helix is present in mammalian NECAP ‘Ex’ segment, with the key binding residues being conserved (see consensus sequence between FCHO2 and Necap1 [LI]K[VL][SC]IGNIT) (Fig.S9B), along with the prediction of helicity (*45*)). Consequently, we believe N3 and Necap Ex domain can bind in the same place on *β*2, which could explain the unassigned density seen in SPA structures of µ2Thr156-phosphorylated AP2 bound to NECAP (*57*) – see Fig.S9B,C,D.

### The FCHO-driven concentration, orientation and activation of AP2 needs to be readily reversible

Since CCV formation is energetically expensive, being able to reverse it i.e. abort CCV formation early (<10s) when optimal conditions or requirements for a CCV’s formation (e.g. PtdIns(4, 5)P_2_ and cargo) are not met (*18, 29, 58*) is important in order to prevent waste of resources. In *in vitro* reconstructions without time constraints, at high concentrations and without any competition for PtdIns(4, 5)P_2_ and cargo (*9, 16*), stochastic membrane encounter is sufficient in the absence of FCHO to form CCPs/CCVs. *In vivo* FCHO absence would be expected to cause a high number of abortive events, with CCPs rapidly disassembled. Indeed, this is observed in cultured cells when FCHO has been deleted (Fig.S8A and (*15*)). However, only a comparatively modest defect results (on average 40-50% reduction in transferrin internalization rates) (*15, 18, 26-29*) - presumably in part due to redundancy built into the system e.g. Eps15/R can still bind AP2 (reviewed in (*59*)). This reduction in CME rate and the resulting energy expenditure can, it seems, be largely tolerated in the energy-rich conditions in cell culture. In whole organisms, although abortive CCV initiation events do naturally occur (*60*), phenotypes of FCHO deletion are marked and penetrative (*12, 61*). Finally, since N1 and N2, which are involved in the PtdIns4,5P_2_-actuated switch on Cµ2, are subject to phosphorylation/dephosphorylation events e.g. by CK2, the system can receive physiological inputs to finely tune the rate of CME and/or the decision as to whether or not to abort or proceed to ordered completion to the needs and condition of the cell.

### Possible roles for the FCHO/Eps15 Corral

Since open AP2 will have coalesced into the centre of the forming CCP under the polymerized clathrin and demonstrates only minimal interaction with open, membrane and cargo bound AP2, FCHO (crosslinked by Eps15) will be excluded from the centre of the forming CCP and displaced to the periphery of the CCP. Thus FCHO/Eps15 will form a ring. If this phase separated structure works similarly to the centre of a nuclear pore (*62*), allowing only proteins that can bind its constituents to cross it (i.e. most CCP components can bind FCHO (*15, 28, 63*) and/or Eps15.(*64*)), then the FCHO/Eps15 ring will bind closed AP2 that is loosely membrane- assocaited on its outside whilst effectively corralling open, fully PtdIns4,5P_2_-bound, clathrin- crosslinked AP2 and to prevent its ‘escape’. Further, any free phosphoinositides and cargo released by clathrin adaptors within the corral would be inhibited from diffusing out of it i.e. it acts as a diffusion barrier. As the number of FCHOs and Eps15 in a CCP plateaus early on in its formation, conversion of the FCHO/Eps15 patch into an annulus of increasing radius but decreasing thickness will only be possible up until the point when it is stretched to its maximum length without breaking i.e. a ring of FCHO dimers linked to each other by Eps15s(*17*). It would then resist further expansion and the resulting maximum diameter of the annulus should help to define the maximum size a CCP/CCV by the number of clathrin adaptors it can corral inside it, providing an explanation for why all CCPs in the RPE and U-2 OS cells are basically of the same size/intensity. It is therefore possible that the amount of FCHO/Eps15 may play a role in defining the standard CCV size/curvature along with the amount of CALM (*37, 65*) Intriguingly an annular structure has also been seen in the endocytic CCPs of yeast (*66*), despite its endocytosis being largely actin dependent and so mechanistically different, suggesting that this design feature is not only conserved throughout eukaryotes but also that its formation may be a common mechanism in many transport vesicle genesis events

To conclude, our eTIRF SIM shows that a small number of FCHO molecules, presumably clustered on the PM by virtue of their FBAR-domains binding PtdIns4,5P_2_ and PS and by forming phase-separated nano clusters with Eps15 ((*17*) (*38*) and Figs. 1D,E) act as potential sites for CCP nucleation. FCHO achieves this primarily by engaging, recruiting and activating AP2 at four sites scattered across AP2’s surface. The apparent low µM K_D_ overall interaction strength of the FCHO•AP2 complex makes AP2 membrane-dwell time, concentration and kinetics commensurate with the physiologically relevant timescales for CME (30-90s)(*10, 39, 59, 67, 68*). However, by being constructed from multiple weak interactions, the formation of FCHO•AP2 complexes is rapidly reversible. As the CCP grows and matures, the FCHO/Eps15 transforms from a patch into a corralling ring surrounding a central, likely domed patch of membrane-attached AP2 and other clathrin adaptors surmounted by a lattice of clathrin itself. However, since the FCHO•AP2 interaction should only occur at their transition zone/interface and stable membrane-attachment of AP2 disrupts FCHO•AP2 complexes, and since AP2s in the central patch will be clustered under a clathrin lattice, released FCHOs will be pushed to the periphery where they can entrap, activate and transfer further AP2s into the central patch as CCP formation progresses.

## Acknowledgements

This work is in memory of Linton Traub, a good man, a good scientist and a good friend who was diagnosed with multiple myeloma during this work’s early stages and battled with this hideous illness throughout most of its length only to sadly die before it could be completed and written up. We also Thank Meir Aridor for helping access and analyse Linton Traub’s data.

We thank the staff and especially the protein crystallography beamline scientists at Diamond Light Source plc. SPA for this study was carried out at the Cryo-EM facility of the University of Cambridge at the Department of Biochemistry (funded by WT 206171/Z/17/Z and WT 202905/Z/16/Z) and we thank Dimitri Chirgadze, Steven Hardwick and Lee Cooper for assistance with sample screening and data collection and Jonathan Wilson for computational support. It also made use of the MRC LMB EM facility and high performance computing resources. NRZ, ZK, VKD, AGW, BTK and DJO were funded by WT 207455/Z/17/Z, ZK was also funded by a WT fellowship 220597/Z/20/Z, VKD also funded by ARUK grant ARUK-PPG2018A-004. LMT and PKU by NIH grant 5R35-GM134855, and PKU also by a Ramalingaswami Fellowship, Department of Biotechnology, India (BT/RLF/Re-entry/18/2014). JAGB and OK by Medical Research Council (MC_UP_1201/16). F.S. and J.K. were supported by Czech Science Foundation (GA18-05360S) and by the Praemium Academiae awarded by the Czech Academy of Sciences. MF and KK thank the Wellcome Trust (212343/Z/18/Z) and EPSRC (EP/S004459/1). The TIRF-SIM microscope was built in collaboration with and with funds from Micron (www.micronoxford.com) supported by Wellcome Strategic Awards (091911 and 107457) and an MRC/EPSRC/BBSRC next-generation imaging award. KK was supported by WT Fellowship (100262/Z/12/Z) awarded to Michael Dustin.

Also, we want to thank Dr Zhibin (Alex) Li and his group (Edinburgh Centre for Robotics), who contributed to early discussions on approaches and tools suitable for eTIRF-SIM data analysis.

## Author contributions

Cell Biology ZK, LMT; TIRF ZK; eTIRF-SIM ZK, KK, MF; Image analysis ZK, JK, FS, KK; Protein production and Molecular Biology NRZ, SG, SH, BTK, DJO, LMT; Crystallographic determinations NRZ, AGW, PRE, DJO; SPA EM VKD, NRZ, DJO, KQ, JB; Tomography OK, JK, JB; Biophysics and biochemistry NRZ, PKU, BTK, DJO, LMT.

Funding ZK, VKD, DJO LMT; All authors contributed appropriately to the design of their experiments and writing of the manuscript. Overall project design and management ZK, BTK, DJO and LMT; Conceptualisation and coordination DJO.

## Data and materials availability

Atomic coordinates and structure factors of AP2-FCHO, Cμ2- FCHO and *α*2 ear -FCHO complexes have been deposited in the PDB under accession codes 7OHO, 7OG1, 7OFP, 7OHZ, 7OI5, 7OIT, 7OIQ and 7OHI, respectively. Crystallographic data collection and refinement statistics are in Tables S3, S4 and S5. Electron density maps and coordinates for the SPA of AP2-FCHO2 chimera have been deposited in the Electron Microscopy Data Bank (EMDB) with ID code XXXX and the Protein Data Bank (PDB) with ID code XXXX, respectively. All other data needed to evaluate the conclusions in the paper are present in the paper and/or the Supplementary Materials. Additional data related to this paper may be requested from the authors.

