## Supplementary Information for "FCHO controls AP2’s critical endocytic roles through a PtdIns4,5P_2_ membrane-dependent switch"

**Short Title:**
**The role of FCHO during CCP formation**

**Authors:**
Nathan R. Zaccai[1*], Zuzana Kadlecova[1*$], Veronica Kane Dickson[1], Kseniya Korobchevskaya[2], Jan Kamenicky[3], Oleksiy Kovtun[4], Perunthottathu K. Umasankar[5], Antoni G. Wrobel[1], Jonathan G.G. Kaufman[1], Sally Gray[1], Kun Qu[4], Philip R. Evans[4], Marco Fritzsche[2,6], Filip Sroubek[3], Stefan Höning [7], John A.G. Briggs[4,8], Bernard T. Kelly[1], David J. Owen[1$], Linton M. Traub R.I.P. [9]

### Supplementary Materials and Methods:

#### Constructs used in this study

PMIB6mScarlet-3xFlagGGGG-FCHO2-WT and PMIB6mScarlet-3xFlagGGGG-FCHO2-STRING mutant : residues 314-444 replaced by GNANGNAGAPAGANGNANPGNAGNGAGAPAGANGNANPGNAGNGANAGNGPGNGANAGANAGNANGNAGAGNANGNANPGAGNGAGNANGNANPGAGNGAGNAGNAGANAGNGPGNGANAGANA

AP2 α−egfp retroviral construct was gift of Schmid lab (*29*)

pEGFP FCHO FL(residues) 1-810 and deletion mutants thereof:
pEGFP FCHO BAR only (residues 1-275), pEGFP FCHO BAR+linker (residues 1-454),
pEGFP FCHO BAR+linkerΔN3 (residues 1-365
+GNANGNAGAPAGANGNANPGNAGNGAGAPAGANGNANPGNAGNGANAGNGPGNGANA+414-454)

AP2 core made with pMWHis$_6$β2trunk+mycμ2, pMWGSTαtrunk+σ2

AP2:μ2FCHO2-N1+N2+N3 chimaeric AP2 core made with pMWGSTαtrunk+σ2 and pMWHis$_6$β2trunk+mycμ2-linker-FCHO2 (314-444)
Linker sequence (GASGSAGSAGPSGAGSAGSAGPSAGSAGSAGSGSAGSAPG)

AP2:β2FCHO2linker made with pMWGSTαtrunk+σ2 and pMWHis$_6$β2trunk(residues 1-542)-FCHO2 (358-444)+mycμ2

pMWHis$_6$μ2 (160-435) wt

pMWHis$_6$μ2 (160-435) – linker - FCHO2 residues (314-351)
Linker sequence (GASGSAGSAGPSGAGSAGSAGPSAGSAGSA)

pGEXμ2FCHO2 (160-435) – linker - FCHO2 residues (314-351)
Linker sequence (GASGSAGSAGPSGAGSAGSAGPSAGSAGSAGSGSAGSAPG)

pETHis$_6$σ2α2 with σ2 residues (1-142) and α2 residues (9-395) (*69*)

His$_6$MBPμ2(160-435) wt and BR4- mutant (K339A/K341A/K343A/K345A)

pGEXFCHO2linker wt residues (314-444) and deletion mutants thereof:

pGEXFCHO2 N1-N2 residues (314-356), pGEXFCHO2 N3-C residues (354-444), pGEXFCHO2N1 residues (314-340), pGEXFCHO2N2 residues (334-356), pGEXFCHO2 N3 residues (354-397), pGEXFCHO2 C residues (413-444)

pGEXFCHO2linkerN1-N3 wt residues (314-394) and mutants thereof EEK (E320S/E321S/K326S), YS1(Y323S), YS2(Y340S) EFAS (E336A/F339S)

pGEXFCHO2 phosphomimetic linker residues (314-444) with mutations (S341E, S342E ,S343E ,S345E ,S347E),

pGEXFCHO1linker wt (N123C) residues (316-444) and deletion mutants thereof: pGEXFCHO1N23C residues (334-444), pGEXFCHO1N13C residues (316-353 and 354-444), pGEXFCHO1N12C residues (316-354 and 414-444), pGEXFCHO1N123 residues (316-413) pGEXFCHO1N3C residues (354-444), pGEXFCHO1C residues (414-444)

pGEX4T2$\alpha$-appendage residues (695-938)

GST SERINC-L10 residues (residues 354-404) gift of Rajendra Singh

pGEX ARRS control (residues (627-635) of $\beta$arrestin2)

pGEX4T2 linker
Linker sequence
(GSGEEVNGDATAGSIPGSTSALLDLSGLDLPPAGTTYPAMPTRPGEQASPEQPSASVSLLDDELMSLGLSDPTPPSGPSLD)

pGEX4T2 linker TGN38
Linker sequence
(GSGEEVNGDATAGSIPGSTSALLDLSGLDLPPAGTTYPAMPTRPGEQASPEQPSASVSLLDDELMSLGLSDPTPPSGPSLDYQRLN)

**Antibodies used**

Affinity-purified rabbit polyclonal anti-FCHO1 antibody 1 (1:2500) (Traub Laboratory),

Affinity purified rabbit polyclonal anti-FCHO2 (1:2500), (Traub Laboratory), NOVUS Biological polyclonal anti FCHO2 NBP2-32694

Affinity purified rabbit polyclonal anti-AP-1/2 β1/β2-subunit GD/1 (1:2500) (Traub Laboratory)

Affinity purified rabbit polyclonal anti-Eps15 (1:1000 HPA008451) (Atlas antibodies)

mAb directed against the AP-2 α subunit clone 8/Adaptin α (1:1000; 610502) (BD Transduction Laboratories, San Jose, CA)

mAb directed against AP-2 μ2 subunit clone 31/AP50 (1:500; 611350) (BD Transduction Laboratories, San Jose, CA).

mAb AP.6 directed against AP2 α subunit gift of Prof Frances Brodsky

Secondary antibodies used were: for blots donkey anti-rabbit (1:5000; NA934V)- or anti-mouse (1:5000; NA931V)- horseradish peroxidase conjugates from GE Healthcare Life Sciences (Pittsburgh, PA): for immunofluorescence goat anti-mouse conjugated to either AlexaFluor488 or AlexaFluor568 (Invitrogen)

### Supplementary Methods

#### Cell Biology

**Cell culture:** Parental U20S cell line was obtained from ATCC. All cell lines were grown under 5%$CO_2$ at 37°C in DMEM high glucose Glutamax medium (Thermo Fisher Scientific) supplemented with 10% (v/v) fetal calf serum (FCS, HyClone). The mycoplasma test was performed routinely with MycoAlert™ PLUS Mycoplasma Detection Kit (Lonza), to rule out mycoplasma contamination.

**Generation of FCHO2 Knockout U-2 OS cell lines by CRISPR-Cas9n:** Genome editing of U-2 OS cells to generate isogenic knockout clones of endogenous FCHO2 (FCHO2-KO) was performed by Genscript. The plasmid containing the human codon optimized SpCas9 gene with 2A-EGFP and the backbone of sgRNA (*70*) was obtained from Addgene. SgRNA CCTGGAGTATGCCTCCTCTA was designed using the CRISPR/Cas9 target online predictor (*71*) to disrupt Exon 2 of FCHO2 (NM_138782.3). The Cas9 plasmid was transfected in the U-2 OS cells and the transfected and FACS sorted eGFP positive cells were plated in 96-well plates by limit dilution to generate isogenic single clones. The clones were expanded and screened by Sanger DNA sequencing to identify positive isogenic single knock-out clones. Three positive knockout clones (#21, #33 and #31) were further validated by RT-PCR and western blotting using NOVUS Biological polyclonal anti FCHO2 NBP2-32694. A wild-type isogenic clone (#6) was maintained as a negative control to generate isogenic pairs of wild-type and mutant cell lines.

**Reconstituting FCHO2 KO cell lines with FCHO2-string and FCHO2-wt constructs:**

The FCHO2 KO cell lines were reconstituted with mScarlet-FCHO2$^{WT}$ or mScarlet-FCHO2$^{STRING}$ using retrovirus-based expression as previously described. The construct termed mScarlet - FCHO2$^{STRING}$ contained a modified linker based on Gly, Asn, Ala, and Pro residues (predicted to have no secondary structure) with an identical length. The different pools of cell lines with mScarlet-FCHO2$^{WT}$ were obtained by FACS into populations of low, medium and high FCHO2 expression, cell pools were further analysed by western blotting. The population with the FCHO2 expression equivalent to that of wild-type U20S cell lines was selected for further experiments. Stable U-2 OS-FCHO2 cell lines carrying fluorescently-tagged CLCa were generated as previously described.

AP2-eGFP cells lines were generated by transducing U-2 OS-FCHO2 cells with the corresponding retrovirus as described (*72*). After 72 hours, cells were isolated in populations with different eGFP expression levels by fluorescence-activated cell sorting (FACS Flow Cytometry, CIMR). Cell pools with an over-expression of eGFP-tagged AP2 subunits at 2x times that of endogenous subunit and at least 70% incorporation were then selected for live- cell imaging experiments.

**Microscopy**

**Live cell TIRF microscopy:** Experiments were performed as previously described. Briefly, high precision #1.5 round coverslips were acid washed and coated with 0.2 mg/mL gelatin (Corning, #2850-22). All TIRF imaging was carried out with a Zeiss 100x 1.49 NA Apo TIRF objective, which was mounted on a Elyra PS1 inverted microscope with a Definite Focus System. Time-lapse series were acquired at a frame rate of 0.5 Hz using a PCO Edge 5.5 sCMOS camera. During imaging, cells were maintained at 37°C in DMEM supplemented with 10% FBS.

**TIRF data analysis:** Single channel and dual channel TIRF movies were analysed with CME analysis suite. Initiation densities of CCPs and short-lived, subthreshold, clathrin structures was determined as previously described (*68, 73*)

**Dual colour TIRF data analysis:** TIRFM movies of each double-labelled cell line expressing either FCHO2-WT or FCHO2-string were obtained. Master/Slave analysis was used to determine the degree of colocalization in bona fide CCPs throughout their lifetimes. In the first analysis AP2-egfp or CLC-egfp was used as a primary channel to determine the evolution of FCHO2 intensity prior to initiation of bona fide CCP. The number of buffer frames was changed from a default value of 5 to 25 frames. This allowed visualizing background and initial FCHO2 intensities in the same pixel before AP2 or egfp-CLCa appearance. This step effectively excluded any CCPs occurring at hot spots from the analysis: A hotspot is defined as a formation of one CCP followed right after by another. Hence these CCPs cannot be used for determining the initial de-novo recruitment of FCHO2. Inflection timepoints of AP2 and FCHO2 averaged intensity cohorts were calculated with an approximating cubic polynomial.

For complementary analytical approach we assigned primary channel to FCHO2 dynamic structures. to determine their capacity to recruit AP2- and egfpCLCa.

**eTIRF-SIM microscopy:** The eTIRF-SIM was performed using a custom-built system described at the Kennedy Institute at the University of Oxford. The detailed description of setup configuration can be found in and applied experimental and analysis pipelines in (*74, 75*). In short, the system consists of a 1.49-NA 100x objective lens (Olympus) fitted on an inverted microscope body (IX-83, Olympus) and complemented with two high-speed sCMOS cameras (ORCA-Flash4.0; Hamamatsu). The key element of the setup is ferroelectric spatial light modulator (SLM) for generating structured illumination pattern and the total internal reflection (TIRF) incidence angles. The SLM switching between different modes at very high repetition rates allows fast imaging with up to 200ms frame rate. To capture CLCegfp and FCHO2 dynamics in live-cell eTIRF-SIM movies 488nm and 560nm wavelengths were used for the excitation. TIRF angles were adjusted for both wavelengths to ensure 100-150nm penetration depth in the axial dimension, which created well confined optical sectioning at the basal plane. Nine raw images (3 angles and 3 phases of structured illumination pattern) for each time point and wavelength were acquired to obtain a super-resolved image with about twice the resolution of the standard TIRF image. The raw images were processed and reconstructed by a previously described algorithm(*76*). The identical Wiener filter parameter of 0.05 was used for reconstruction of all the images.

**eTIRF-SIM image analysis**

**Pre-processing:** Prior to image analysis images were chromatically corrected using Multistackreg macro in Fiji (Schindelin et al., 2012). To ensure high quality correction, the multi-color images of 100nm Tetraspeck microspheres (Thermo fisher, T7279) images were taken before each data set and used as transformation matrixes.

**Automatic Detection and Tracking:** We adopted CMEanalysis software (*68, 73*), which is an excellent tool for detecting and tracking CCPs in TIRF time-lapse movies. However, in its original version it is not suitable for reconstructed eTIRF-SIM images with annular CCPs, since it models fluorescent signal of a CCP as a two-dimensional Gaussian approximation of the microscope PSF above a spatially varying local background. Hence, we averaged the raw 9 images acquired for each time point and each channel to generate a single snapshot per channel and time point for CCP Detection and tracking in CME analysis with default parameters. The resulting detected spatial coordinates of valid tracks were superimposed on the final eTIRF-SIM reconstructed image. This approximate location and track for each CCP was then manually processed, and each ROI was scrutinized and manually CCP centroids were re-positioned automatically in both channels if necessary. For this purpose, we developed a simple tool with an intuitive GUI to speed-up this step. Only single CCPs with canonical lifetime profile were adjusted and used for downstream analysis.

**Manual post-processing and intensity analysis, averaging, bleaching correction:** We then manually refined the position of the detection in respect to CCP centroids in eTIRF-SIM data, since the tracks obtained with CME analysis from raw TIRF images only approximated it.

**Bleaching correction:** When extracting individual CCP regions from reconstructed eTIRF-SIM data, we need to compensate for intensity bleaching. We do this by averaging intensities in individual time frames and then fit an exponential curve of the form Y=a*exp(b*x)+c*exp(d*x) to this data which gives us the final bleaching compensation coefficients.

**Intensity adjustment and averaging:** We preserve original intensities in individual channels. When comparing different channels, we estimate a multiplicative coefficient, which minimizes the difference between average histograms of the channels as a least squares fit.
Finally, at least 50 snapshots defined as 21x21 pixel ROI for a given phase of individual CCPs were assembled in a stack and averaged using Z-projection and intensity averaging in Image J.

**Protein expression and purification**

Recombinant proteins were expressed in BL21 plyS *E. coli* grown in 2TY media. After $OD_{600} > 0.6$ cell density was reached at 37°C, expression was induced with 0.2mM IPTG overnight, shaking at 22°C. Cells were lysed using a cell disruptor (Constant Systems). AP2 core and AP2 chimeras were made in 250mM NaCl 10mM Tris pH 8.7 2mM DTT according to previous protocols (*14*). Briefly, the complexes were initially isolated with GST-beads and cleaved overnight with thrombin. The resultant protein was then isolated on NiNTA-beads, washed with buffer supplemented with 10mM imidazole, and eluted from the beads with the same buffer supplemented with 300mM imidazole. After size exclusion chromatography on a Superdex 200 column (GE Healthcare) in 250mM NaCl 10mM Tris pH 8.7 2mM DTT, the AP-2 complexes were concentrated to >15mg/ml with vivaspin concentrators.

Recombinant Cμ2 and other $His_6$-tagged proteins were made as in (*47*). Briefly, the proteins were prepared in 500mM NaCl 20mM Tris pH 8.0. Proteins were initially isolated on NiNTA agarose, washed with buffer containing 20mM imidazole and eluted with buffer supplemented with 300mM imidazole. GST-tagged Cμ2 and α-appendage proteins were made as in (*51*). Briefly, the proteins were prepared in 200mM NaCl 20mM Tris pH 7.4 1mM DTT. Proteins were initially isolated on GST beads, washed and cleaved from their tags over night at room temperature with thrombin. The final purification step for both $His_6$-tagged and GST-cleaved proteins involved size exclusion chromatography with a superdex 200 column (GE Healthcare) in their respective preparation buffers or in the case of material for ITC into 150mM NaCl 100mM Tris pH 7.4 1mM DTT. Proteins were subsequently concentrated to >10mg/ml with vivaspin concentrators. GST FCHO linkers were similarly purified on GST Sepharose with overnight thrombin cleavage where necessary and finally by S200 gel filtration with all purification done in HKT buffer (10mM HEPES, 10mM Tris pH 7.4, 120mM potassium acetate, 2mM DTT)

**Crystallography**

**Crystallization:** Crystals of AP-2 FCHO2 chimera were grown in hanging drops with reservoir 18% PEG 12000 0.1M Na/K phosphate pH 6.2 0.2M NaCl 4mM DTT in the presence of 3-fold molar excess of IP6. Crystals were cryo-protected with 20% glycerol.

Apo crystals of recombinant histidine-tagged Cμ2 were grown in sitting drops from a mixture of Cμ2 (10 mg/ml) and the FCHO2-derived N1 block peptide (DVDEEGYSIKPETNQNDTKENHFYSS) (2mg/ml) equilibrating against a reservoir containing 1.5M Ammonium Sulphate 0.1M HEPES pH

7.0. Crystals were cryo-protected by soaking in mother liquor supplemented with 20% glycerol and peptide. The N1 peptide was not visible but Trp421 had swung round to partly fill the Y-binding pocket to reduce its solvent exposure (Fig.S6 D,E,F)

Crystals of recombinant histidine-tagged Cμ2 in complex with the FCHO2-derived C block peptide (SDLLAWDPLFG) were grown in sitting drops equilibrating against a reservoir containing 20mM sodium formate; 20mM ammonium acetate; 20mM sodium citrate tribasic dihydrate; 20mM sodium potassium tartrate tetrahydrate; 20mM sodium oxamate, imidazole MES monohydrate pH 6.5 , 20% v/v glycerol; 10% w/v peg 4000 ($P3_221$ crystal form) and in 30mM magnesium chloride hexahydrate, 30mM calcium chloride dihydrate, 100mM sodium HEPES MOPS pH 7.5, 20% v/v ethylene glycol; 10 % w/v peg 8000 (C2 crystal form). The crystals did not require cryo-protection.

Recombinant histidine-tagged Cμ2-FCHO2 chimera crystals were grown in sitting drops against a reservoir containing 20% w/v PEG 3,350 0.2 M DL-Malic acid pH 7.0. The crystals were cryo-protected by soaking in mother liquor supplemented with 30-32% glycerol.

Recombinant GST-cleaved Cμ2-FCHO2 chimera crystals were grown in sitting drops against a reservoir containing 20% w/v PEG 3350, 0.2 M Sodium phosphate dibasic dehydrate pH 9.1 and were cryo-protected by soaking in mother liquor supplemented with 25% glycerol.

Crystals of AP-2 in complex with FCHO2 linker and the DYQRLN peptide derived from TGN38, supplemented with 10mM K Na Tartrate, grew in sitting drops with reservoir 0.1M Magnesium formate dehydrate, 10% to 15% PEG 3350. The crystals were cryo-protected with 0.1M Mg formate dehydrate, 13% PEG 3350, 18-24% Glycerol and 1mg/ml of peptide. Crystals of AP-2 in complex with selenomethionine-labelled FCHO2 linker (made as in (*16*)) and the DYQRLN peptide were grown in similar conditions.

Crystals of α2 ear complexes were grown in sitting drops from a mixture of α2 ear (10 mg/ml) and the FCHO1-derived C block peptide (QSEEQVSKNLFGPPLESAFDHED) (2mg/ml) against a reservoir containing 1.0M Lithium sulphate 0.1M MES pH6.5. Crystals were cryo-protected by soaking in mother liquor supplemented with 20% glycerol and peptide.

**Synchrotron data collection and structure determinations:** Diffraction data were collected at Diamond Light Source on beam lines I03, I04 and I24 and data processed with Xia2. Initial structures were solved by molecular replacement with the program PHASER by using as search models the previously published structures of Cμ2 (PDB 1BXX), α-appendage (PDB 1W80) and AP2 (PDB 2VGL and 2XA7). The positions of the selenomethionines in the structure of AP-2 in complex with TGN and Selenomethionine-labelled FCHO2 were determined from the anomalous diffraction collected at wavelength 0.92-Angstrom. Iterative rounds of refinement with the programs PHENIX REFINE and REFMAC were interspersed by manual rebuilding of the model with COOT Crystallographic programs were run from the PHENIX (*77*)) and CCP4 packages (*78*)) and figures were produced with a commercial version of Pymol (Schrödinger, LLC). Data collection and refinement statistics are in Tables S3, S4 and S5.

**Single Particle Cryo Electron Microscopy**

AP2-FCHO2 chimera purified as wt AP2 and finally placed in in 50% HKT buffer (10mM Hepes, 10mM Tris 120mM potassium acetate pH 7.2) and 50% Core buffer (10mM Tris, 250mM NaCl, pH

8,7) was applied to Quantifoil grids of type R1.2/1.3 on 300 copper mesh +/- 0.05% ß-octyl glucoside. Grids were glow-discharged for 60 seconds at 20 mAmp using a Pelco EasiGlow before application of 3.5 µL of sample (0.4 mg/mL) and plunge-frozen using a Vitrobot Mark IV (FEI Company) operated at 4°C and 95% humidity. Data collection was carried out on a Titan Krios transmission electron microscope (FEI/Thermo) operated at 300 keV, equipped with a Gatan K3 direct detector (FEI) in counting mode. Automated data acquisition was performed using FEI EPU software at a nominal magnification of 130,000, which corresponds to a pixel size of 0.362 Å per pixel in super resolution mode. Dose-fractionated movies were acquired using 1.31 second exposures and 48 fractions at a dose of 36.09 e/Å2/sec in the defocus range of -0.8 to -2.8 µm.

Data collection quality, movie frame alignment, estimation of contrast-transfer function parameters, particle picking and extraction were carried out using Warp (*79*). 718,511 particle images were extracted with a box size of 340 and imported into CryoSPARC(*80*) for particle curation. 2D classification, ab initio model building, 3D refinement, filtering and sharpening was also carried out using CryoSPARC as described in the workflow (Fig. S5). After 2D classification, 372,423 particles were retained in the dataset which was subjected to further 3D classification steps. After the first round of 3D refinement, particles assigned to a poorly resolved class, and to the CMu-out class, were removed from the dataset. The remaining particles were further refined to generate the CMu-in structure and subject to further 3D classification to generate the N1-N2 enriched structure. Analysis of particle orientation distribution, and assessment of anisotropic resolution by calculating the FSC within cones of 30 degrees in 3DFSC (81) showed that for both structures the resolution in the Y direction was lower than in the other two directions. Random particle subsets in the oversampled views were removed from the dataset to reduce the resolution anisotropy and retained particles were further refined to reconstruct the final map. Structures were filtered according to local resolution using cryoSPARC for visualization and interpretation.

The previously-determined 'closed' conformation of the AP2 core complex (PDB ID 2vgl) was fitted into cryo-EM volumes using the phenix.dock_in_map function of the Phenix Software suite (CC values reported in Table S5). The structures were refined using phenix.real_space_refine (*81*).

**Cryo electron tomography**

**Liposome preparation**: The required mixtures of lipids were assembled in 4:1 chlorophorm/methanol, dried and rehydrated in HKT buffer, subjected to five freeze-thaw cycles and extruded eleven times through carbon membranes with 100 nm pores.

**Cryo-electron tomography sample preparation and data acquisition:** AP2 was recruited to liposomes alone or in the presence of the FCHo2 linker by mixing 1.5 µM of AP2, or 1.5µM of AP2 and 7.5 µM of FCHO2 linker, with 0.2 mg/ml of 100 nm extruded liposomes containing 10% brain PtdIns(4,5)$P_2$ and 10% DOPS in a POPC/POPE (3:2) mixture in HKT buffer. Reactions were performed in parallel and incubated for 30 minutes at 21°C. Post-incubation, the reactions were supplemented with 1:10 of 10 nm gold fiducial markers in HKT buffer. 3.5 µl of this mixture was applied on a glow-discharged holey carbon grid (CF-2/1-3C, Protochips) and back-side blotted for 4 seconds at relative humidity 98% and 18°C followed by plunge-freezing in liquid ethane (Leica EM GP2 automatic plunger).

Data acquisition was performed as in (*9*). Dose-symmetrical tomographic tilt series (*82*) were collected in FEI Titan Krios electron microscope operated at 300 kV with a Gatan Quantum energy filter with a slit width of 20eV and a K3 direct detector operated in counting mode using tilt series controller in Serial EM software. Tilt series contained 41 tilted images (-60° to +60° with 3° increment) with 10-frame movies acquired for each tilt and were imaged with a total exposure of ~130 $e^-/A^2$ equally distributed between tilts. The details of data collection are given in Table S1.

**Image pre-processing and tomogram reconstruction:** Movie frames were gain-corrected, aligned and integrated into individual tilt images using align frames (IMOD package) (*83*). Several tilt images and some entire stacks were discarded due to tracking errors during acquisition (Table S1). The tilt images were low-pass filtered according to accumulated dose (*84*), aligned using fiducial markers, and 4-times binned tomograms were reconstructed by weighted back-projection in Etomo (IMOD) for particle picking purposes. For 3D contrast transfer function (CTF) corrected tomograms, defocii and astigmatism of individual non-dose filtered tilts were estimated in CTFPLOTTER (IMOD) (*85*) and phase-flipping CTF-correction and tomographic reconstruction was done in novaCTF (*86*) using 15 nm strip width. Tomograms were binned by 2, 4 and 8 times (hereafter called bin2, bin4 and bin8 tomograms) with anti-aliasing.

**Subtomogram alignment:** Subtomogram alignment, averaging and classification were done as previously described in (*9*) in subTOM packages

https://www2.mrc-lmb.cam.ac.uk/groups/briggs/resources

https://github.com/DustinMorado/subTOM/releases/tag/v1.1.4.

Dynamo (*87*) and Relion (*88*) packages were used for mask making.

**Picking initial subtomograms and alignment to the membrane:** Initial position were picked as described previously (*9*) using the "Pick particle" Chimera plug-in (*89*), approximating liposomes by spheres with uniform surface sampling at every 42 Å. The particles were oriented normally to the sphere surface with random in-plane angle. Subtomograms were extracted at these geometrically-defined positions and orientations from bin8 tomograms and averaged producing initial models. The membrane alignment of subtomogram were then refined by allowing only shifts normal to the membrane and a conical angular search range of ±30°. Particles that failed to align to the membrane were removed based on a cross-correlation threshold selected manually for each tomogram.

**Subtomogram averaging of AP2**: The previously published EM map of AP2 on tyrosine-cargo containing membranes (*9*) was low-pass filtered to 42 Å and used as a reference to find initial AP2 positions. Starting from the initial positions and orientations determined by membrane alignment, bin4 subtomograms were aligned to this reference with lateral shift limited to 80 Å to confine the alignment within the area occupied by a single AP2. Where subtomograms had converged to positions within 70 Å of one another, we selected the subtomogram with the highest cross-correlation score and discarded all others. Subtomograms were further subjected to classification in the subTOM package to remove "empty" subtomograms. The remaining subtomograms were then sorted into identically sized odd and even subsets by vesicle and iterative subtomogram alignment and averaging was performed in bin2 and then in bin1, gradually decreasing the search space and increments for angular and spatial parameters, and moving the low pass-filter moved towards higher resolution. Upon alignment convergence in bin2, particles that had diverged from membrane alignment were removed using a cross-correlation threshold defined manually for each tomogram. Table S2 shows data processing statistics.

### Brain cytosol and HeLa cell extract preparation

Rat brain cytosol was prepared in a homogenization buffer of 25 mM Hepes-KOH, pH 7.2, 250 mM sucrose, 2 mM EDTA and 2 mM EGTA. Rapidly thawed frozen brain tissue was homogenized in a Waring Blender at 4°C in the presence of homogenization buffer with 2 mM PMSF, 5 mM benzamidine and complete protease inhibitor tablets. The resulting thick homogenate was centrifuged at 15,000 *g* for 20 min and the postnuclear supernatant fraction recentrifuged at 17,500 *g* for 20 min. The supernatant was then centrifuged at 105,000 *g* for 60 min. The resulting high-speed supernatant (cytosol) was stored in small frozen aliquots at -80°C.

HeLa cell Triton X-100 lysates were prepared by first collecting confluent cells from Petri dishes using Cellstripper and centrifuging at 500 *g* for 5 min. The cell pellet was resuspended in cold 25 mM Hepes-KOH, pH 7.2, 125 mM potassium acetate, 5 mM magnesium acetate, 2 mM EDTA, 2 mM EGTA, 2 mM DTT and 1% Triton X-100 and incubated on ice for 30 min with occasional mixing. After centrifugation at 24,000 *g* for 20 min to remove insoluble material, the supernatant (lysate) was stored in frozen aliquots at - 80°C. Thawed cytosol/lysate samples for assays were centrifuged at 125,000 *g* for 20 min at 4°C immediately before use.

### GST pull-down assays

A measured amount of GST or GST-fusion protein was immobilised onto 60µl of 50% slurry of glutathione-Sepharose beads in microfuge tubes and the volume made up to 750ul with buffer. After incubation at 4°C with continuous mixing, the Sepharose beads were recovered by centrifugation (10,000 *g*, 1 min) and the supernatants removed. Each bead pellet was washed three times with cold assay buffer and the majority of the buffer was carefully aspirated, leaving a final equivalent volume of ~50µl.

Using cytosols: Aliquots of rat brain cytosol or HeLa cell Triton X-100 lysate were thawed and centrifuged at 125,000 *g* for 20 minutes at 4°C to remove insoluble particulate matter before addition of 200µl to the tubes of immobilized GST or GST-fusion protein and then incubated at 4°C for 60 min with continual mixing. Binding assays were terminated by centrifugation (10,000 *g*, 1 min, 4°C) and 60ul removed and transferred to a new microfuge tube. The pellets were then washed by centrifugation three times with 1 ml/wash ice-cold buffer and the supernatants aspirated and discarded. The Sepharose beads were resuspended in reducing SDS sample buffer, and the volumes adjusted manually so that all tubes were equivalent. After boiling and centrifugation (12,000 *g*, 1 min), aliquots of samples were fractionated by SDS-PAGE and stained either with Coomassie brilliant blue or blotted with indicated antibodies.

**Isothermal titration microcalorimetry**

Experiments were performed using a Nano ITC machine from TA Instruments. For ITC proteins were gel filtered into 100 mM Tris pH 7.4, 150mM NaCl, 0.25 mM TCEP. Peptides were dissolved in the same buffer. PHear domains at concentrations between 0.10 and 0.15 M were placed in the cell at 12°C and peptides at concentrations between 1 and 5 mM (depending on a peptide) were titrated in with 20 injections of 2.43 µl each separated by 2.5 minutes. A relevant peptide-into-buffer blank was subtracted from all data and for constructs, which displayed measurable binding, a minimum of three independent runs that showed clear saturation of binding were used to calculate the mean $K_D$ of the reaction, its stoichiometry (n), and their corresponding SEM values. Analysis of results and final figures were carried out using the NanoAnalyzeTM Software.

**Bio-layer interferometry experiments**

Real-time kinetic measurements of AP2 core binding to immobilized GST-FCHO1linker and GST-FCHO2linker at 25 °C were conducted using an Octet RED96 (Pall FortéBio). AP2 concentrations varied by two-fold dilution series (13.5 – 0.8 µM and 10.0-0.2 µM respectively). Samples and buffers were dispensed into 96-well microtiter plates (Greiner) at a volume of 200 µl per well. Anti-GST biosensors (Pall FortéBio) were incubated in the specific assay buffer for at least 30 minutes prior to loading with the appropriate GST-FCHO linker for 5 minutes, followed by blocking with GST-ARRS control for 10 minutes. Negative controls were analysed with biosensors uniquely loaded with the GST-ARRS control. Every binding experiment consisted of three stEps: incubation for 10 minutes in the specific assay buffer, followed by 15 min incubation with AP2 in the same buffer (association phase), and then by 10 min incubation with the same buffer without AP2 in order to measure AP2 off-rate (dissociation phase). Non-specific binding was reduced by supplementing buffers with 0.1% NP40 and 1mg/ml BSA.

A similar experimental set-up was used in the analysis of AP2 core and (AP2:µ2FCHO2N1+N2+N3) chimera binding to immobilized GST-TGN. The negative control was GST-Control. Binding were measured for the same concentrations in parallel against both the TGN38-loaded and the Control only-loaded biosensors. Two-fold dilution series of AP2 core and AP2 FCHO2 chimera were analysed (3.3 – 0.1 μM and 2.6 – 0.9 μM respectively). Due to slower on- and off-rates, association times were increased to 20 minutes, and dissociation times to 25 minutes.

Data were processed with the Octet Data Acquisition 7.0 software, individually fitted using ForteBio Analysis Software 7.0, and subsequently further analysed with the PRISM software (GraphPad) in order to determine equilibrium dissociation constants ($K_D$). The maximum binding capacity of each biosensor was scaled with respect to each other, according to the molecular weight of the AP2 analysed.

**Liposome spin downs**

GST-FCHO2 WT linker or a non-binding control (GST-ARRS control) at a final concentration of 30 μM was incubated at 4°C for 30 minutes with AP2 core at a concentration of 2.5μM, in 95% HKT buffer with 0.2mg/ml lysozyme as a 'carrier', AEBSF (25 μg/ml) and 5mM DTT. Core buffer (contributed by the purified AP2) was restricted to 5% of the volume. The mixture was then centrifuged briefly to remove any insoluble material. Samples were removed and added to either an equal volume of liposomes (containing PC/PE, PC/PE/PS/PtdIns(4,5)$P_2$ or PC/PE/PS/PtdIns(4,5)$P_2$/TGN38) or an equal bed volume of GSH sepharose beads pre-washed into HKT buffer. Final concentrations were therefore 15μM (GST-FCHO2/ARRS) and 1.25μM AP2. The mixtures were incubated for 30 minutes at 21°C with continuous gentle inversion, then the liposomes were pelleted by centrifugation, supernatants removed and the pellets resuspended in an equal volume before the addition of SDS loading dye and analysis by SDS-PAGE. The GSH-sepharose beads were washed three times with 1 ml HKT supplemented with 0.2 mg/ml lysozyme, AEBSF (25 µg/ml) and 5mM DTT, then resuspended in the same buffer and analyzed by SDS-PAGE. All experiments were done in triplicate.

**Supplementary Figures and legends contents**

**Fig.S1 Validation of FCHO2 levels in U-2 OS cells.**

**Fig.S2 Live cell TIRF microscopy in engineered U-2 OS cells.**

**Fig.S3 AP2 contains overlapping binding sites for FCHO linker and PtdIns4,5$P_2$.**

**Fig.S4 FCHO2 linker can bind intramolecularly to closed AP2.**

**Fig.S5 Single particle cryo-EM image processing workflow.**

**Fig.S6 Structures of Cμ2:FCHO2-N1+N2 chimaeras.**

**Fig.S7 FCHO linker binds to [illegible]2 and μ2 subunits of AP2 using blocks N3 and N1 respectively**

**Fig.S8 Details of FCHO C block binding to [illegible]-appendage and to Cμ2**

**Fig.S9 FCHO linker binding to full length AP2:cellular effect and molecular model**

**Fig.S10 Competing interactions for FCHO linker blocks on AP2**

**Table S1. Cryo-electron tomography data collection.**

**Table S2. Subtomogram averaging image processing parameters and statistics.**

**Table S3 Crystallographic data collection and refinement statistics (AP2/FCHO)**

**Table S4 Crystallographic data collection and refinement statistics (Cμ2/FCHO)**

**Table S5 Crystallographic data collection and refinement statistics (α-ear/FCHO)**

**Table S6. Cryo-electron microscopy data collection, refinement and validation statistics for single-particle structures of AP2:β2FCHO2linker chimaera in solution.**

**Video S1-S4 Super-resolution live-cell imaging of CME dynamics by eTIRF-SIM in U-2 OS cells**

**Video S5 Conformational variability in single particle EM reconstruction of AP2 FCHO2 chimera.**

Zaccai, Kadlecova Figure S1

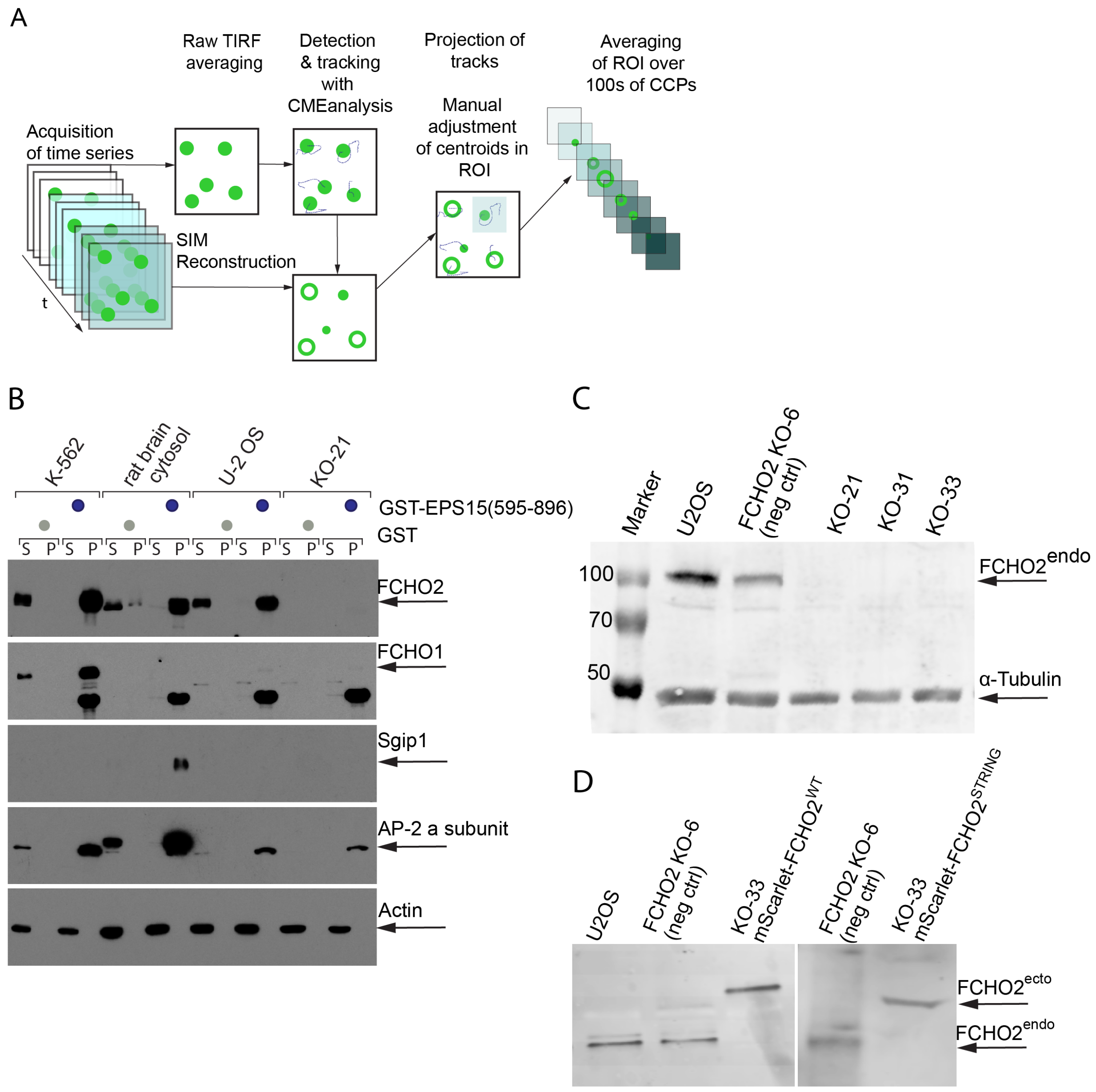

**Fig.S1 Validation of FCHO2 levels in U-2 OS cells.**

**A** Simplified scheme of the analytical workflow used for characterizing CCP formation in live cell eTIRF-SIM movies: U-2 OS cells expressing fluorescently tagged mScarlet-FCHO2$^{WT}$ and egfp-CLCa at near endogenous levels were imaged with eTIRF-SIM in which nine raw TIRF images in the 9 orientations of the grid pattern are acquired to produce a reconstructed super-resolved TIRF-SIM image with about 110nm resolution in both channels. The detection and tracking were obtained for individual CCPs with CMEAnalysis package from the averaged raw TIRF images prior to reconstruction. The centroids of CCPs and tracks were manually aligned simultaneously in both channels to allow ROI extraction and downstream analysis (see M&M). These ROI were averaged for individual CCPs and lifetime phases of CCP formation.

**B** Western blot analysis of the protein quantity of the three FCHO paralogues (muniscins): FCHO1, FCHO2 and Sgip in U-2 OS, U-2 OS FCHO2 KO clone 21, K-562 and rat brain cytosol.

**C** Validation of isogenic U-2 OS FCHO2 knockout clones 21, 31 and 33 and negative control clone 6 by western blot.

**D** Western blot validation of FCHO2 reconstitution in FCHO2 knockout clone 33 by retrovirus mediated expression of mScarlet-FCHO2$^{WT}$ or mScarlet-FCHO2$^{STRING}$ at near-endogeneous levels.

Zaccai, Kadlecova Fig.S2

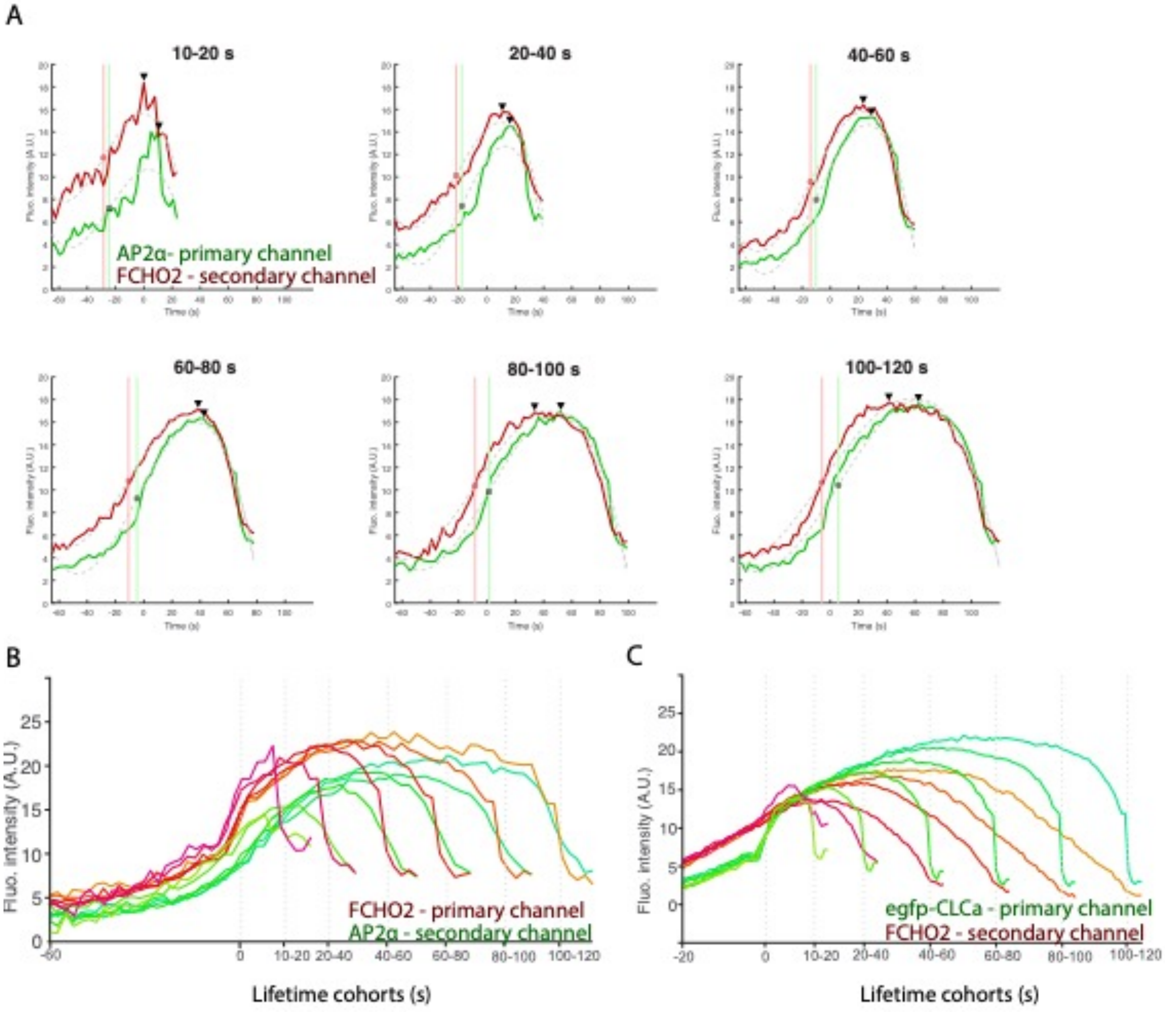

**Fig.S2 Live cell TIRF microscopy in engineered U-2 OS cells.**

**A** Average AP2 (green) and FCHO2 (red) fluorescence intensity traces in lifetime cohorts of FCHO2-positive CCPs. Average fluorescence intensity was plotted for 25 frames prior to the first detected time point in the reference AP2 channel. This allowed visualizing initial FCHO2 intensities in the same pixel before AP2 or egfp-CLCa appearance. Black triangles indicate maximum intensity time point. Red circle indicates inflection timepoints based on cubic polynomial approximation of cohorts (grey dashed lines) for FCHO2 trace. Green circle indicates inflection time point for AP2.

**B** Average FCHO2 (red tones) and AP2 (green tones) fluorescence intensity traces in CCP lifetime cohorts. In this analysis FCHO2 was chosen as the reference channel to define the intensity time courses of AP2 at the time points and locations defined by FCHO2 signal detection. 96% of all dynamic FCHO2 assemblies were classified as positive for AP2.

**C** Average FCHO2 (red tones) and egfp-CLCa (green tones) fluorescence intensity traces in CCP lifetime cohorts.

Zaccai, Kadlecova Figure S3

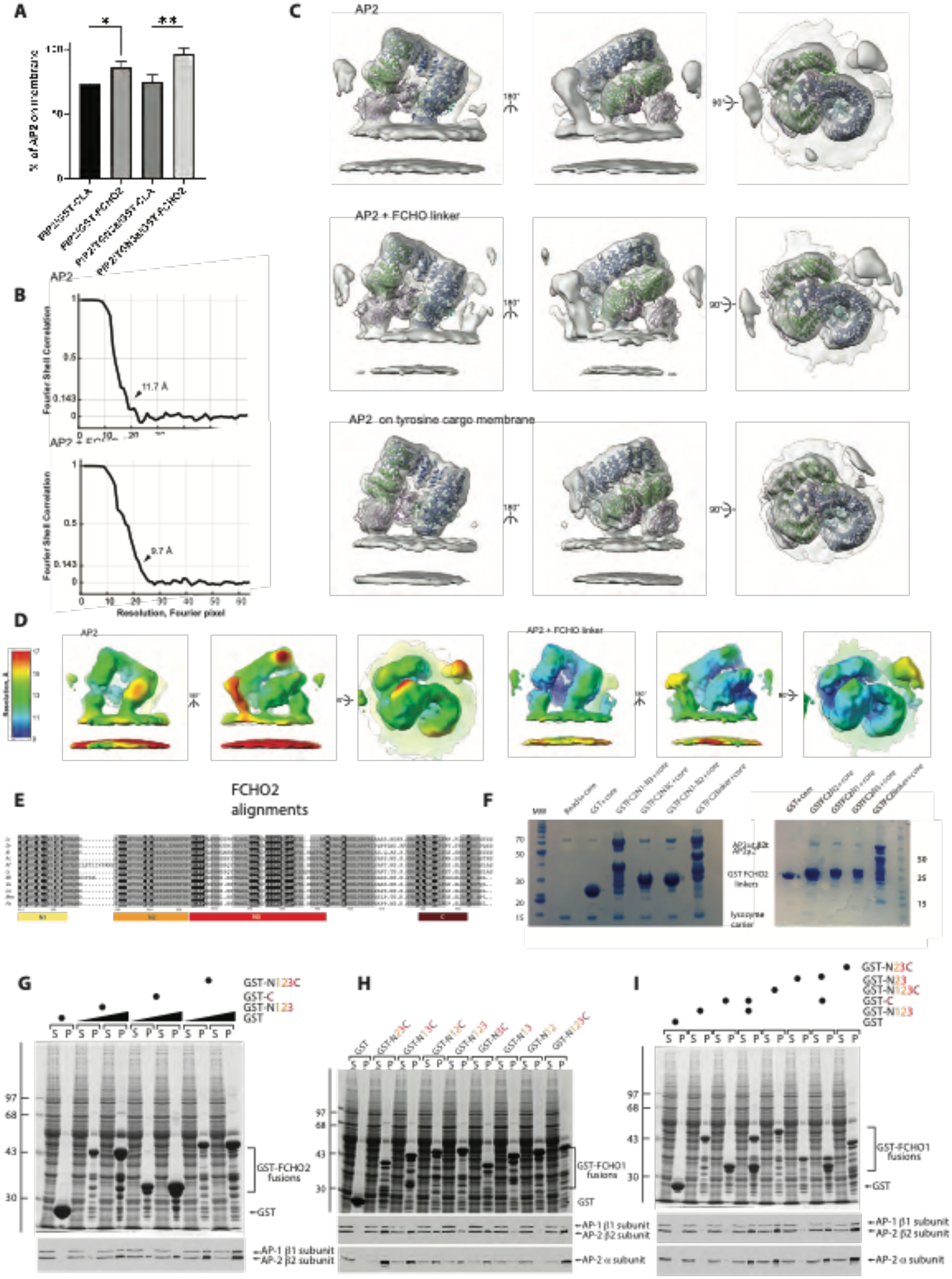

**Fig.S3 AP2 contains overlapping binding sites for FCHO linker and PtdIns4,5$P_2$**

**A** Comparison of AP-2 recruitment to liposomes in the presence and absence of GST-FCHO2. Liposomes (either PC/PE/PtdIns(4,5)$P_2$/PtdSer or PC/PE/PtdIns(4,5)$P_2$/PtdSer/YXXØ cargo peptide) were incubated with mixtures of GST-FCHO2 or GST-ARRS (a non-binding control) and AP-2, the liposomes were pelleted by centrifugation and protein present in the pelleted and supernatant fractions analyzed by SDS-PAGE. The percentages of AP-2 recruited to each type of liposome (assessed by densitometry) are shown as means $\pm$ standard error on the mean (3 independent experiments). By Student's t-test, the percentage of AP-2 recruited was increased by the presence of GST-FCHO2 for both liposome types (PtdIns(4,5)$P_2$, $p = 0.023$; PtdIns(4,5)$P_2$/ YXXØ cargo, $p = 0.006$).

**B** Comparison the conformations of membrane-recruited AP2. Cryo-EM maps for AP2 on membrane without cargo in the absence or presence of five-fold molar excess of the FCHO2 linker, and the previously published structure of AP2 bound to YxxΦ motif containing membranes in the absence of FCHO2 (EMDB-10748). The cryo-EM maps are filtered to 13 Å and all are fitted with the ribbon model of AP2 on the YxxΦ motif containing membranes (PDB: 6YAF). In all cases AP2 is recruited in its open state and there are no observable differences in the conformations at the given resolution.

**C,D** Global and local resolution of on-membrane AP2 EM maps. **C**, EM maps colored by local resolution for both structures determined in this study and **D**, corresponding FSC plots with arrows indicating the measured global resolution at the 0.143 threshold.

**E** Alignment of FCHO2 linker region from species spanning ~400 million years with most distantly related species to humans shown at top. Coelacanth Lc; Zebrafish Dr fcho2; Xenopus Xt fcho2;

Anole Chameleon Ac fcho2; Emperor penguin Af; Society finch Ls; Brown bat Ml; Wombat Vu; Elephant La; Mouse Mm; Human Hs. Identities are shown in black and similarities in grey. The coloured boxes denote the N1, N2, N3 and C block definitions as defined in the human FCHO2 protein and are used throughout the work.

**F** Binding of constructs of GSTFCHO2 linkers containing conserved sequence blocks (as in Fig. 2E and S3E) to recombinant AP2 cores. 30µg of GST or GST-FCHO2 linker fusion proteins as indicated were immobilised onto 60µl of 50% slurry of glutathione-Sepharose beads. Binding was carried 750ul for 30minutes at 4°C with continuous mixing. Sepharose beads were washed 3x 1ml buffer and run on a 12.5% SDS PAGE gel with MW markers. The gel was stained with Coomassie blue: data is summarised in Figure 3F. Removal of C block has little effect on AP2 core binding. Removal of any further block causes a dramatic reduction in binding. Single blocks bind only very weakly.

**G** Binding of 10µg (left) and 30µg (right) of the indicated GSTFCHO2 linkers (Fig. 2E and S3E) to AP2 contained in brain cytosol as in **F** and detailed in Methods: Upper panel Coomassie staining of SDS PAGE gels and lower panel blotting with anti $\beta1/\beta2$ antibody. S soluble fraction. P pellet fraction. Data is summarised in Fig. 3F deletion of C block reduces binding to AP2 when compared with full length linker and C block retains some AP2 binding.

**H, I** Binding of GSTFCHO1 linker constructs containing conserved sequence blocks as indicated in Fig. 2E to AP2 derived from cell cytosol. (**H**) uses single constructs and (**I**) uses combinations of the same as indicated. Binding experiments were carried out as in **G** and outlined in Methods Upper panels Coomassie staining of SDS PAGE gels; middle and lower panels blotted with anti $\beta1/\beta2$ antibody and anti $\alpha$ antibody. S soluble fraction. P pellet fraction.

**H** Deletion of any one block has minor effects on the binding of the FCHO1 linker whereas deletion of more than one block severely compromises binding**:** All blocks show some very weak binding to intact brain cytosol derived AP2. **I** Combinations of various sequence blocks (N1+N2+N3) and C or (N2+N3) and C are more than additive for binding to the individual block constructs indicating cooperativity i.e. avidity effects are occurring

Zaccai, Kadlecova Figure S4

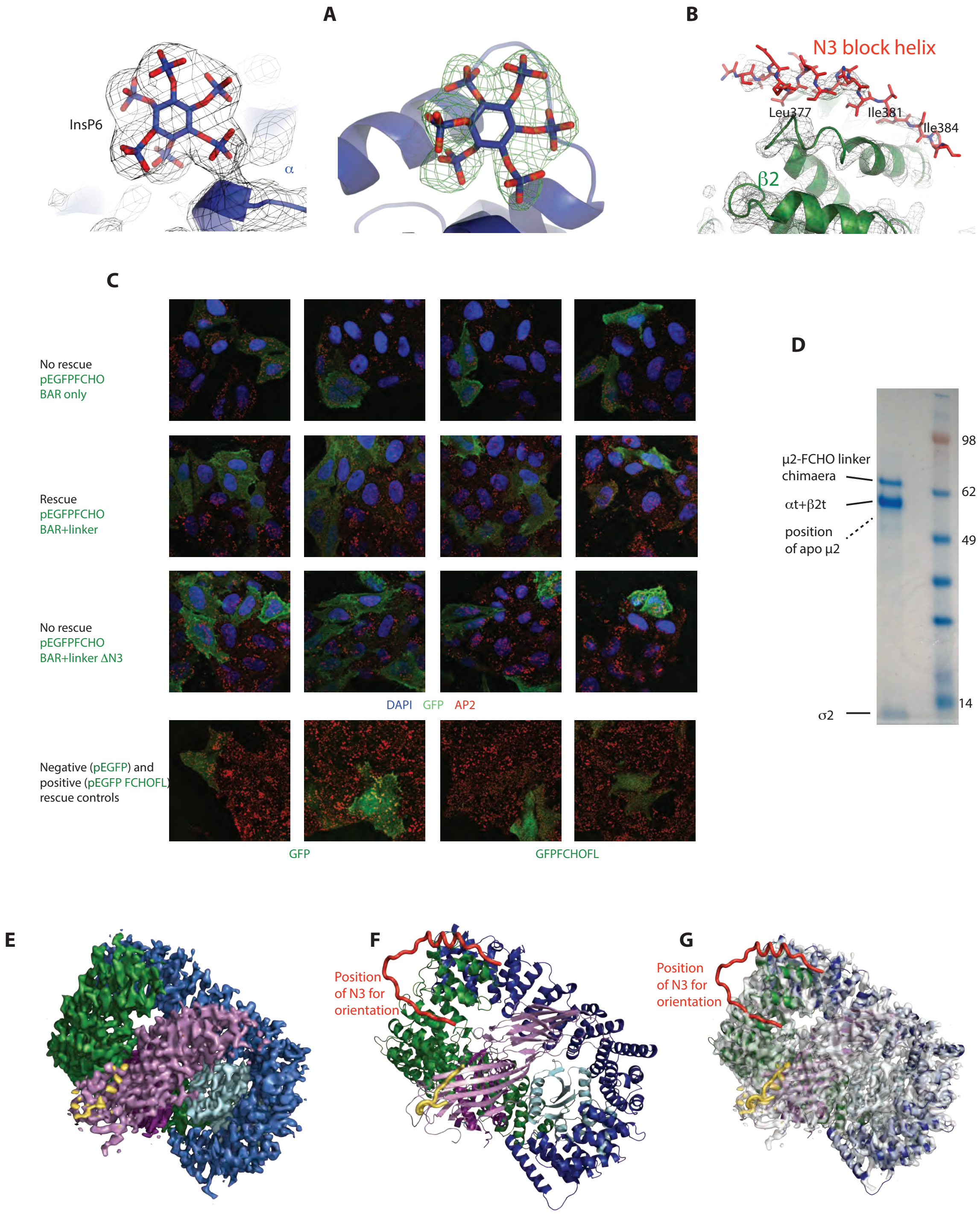

**Fig.S4 FCHO2 linker can bind intramolecularly to closed AP2.**

**A** Left hand panel 2.9Å resolution 2Fo-Fc electron density of D-myo-inositol-1,2,3,4,5,6-hexakisphosphate (InsP6; PtdIns4,5P2 analogue) bound to the [illegible] subunit (blue) of AP2/FCHO chimaera in closed conformation contoured at 1σ. During molecular replacement, InsP6 was not included in the search model. Right hand panel – Fo-Fc omit map for InsP6 contoured at 3σ (green electron density) calculated after simulated annealing with the InsP6 occupancy set to zero.

**B** 2.9Å resolution 2Fo-Fc electron density contoured at 1σ of the AP2/FCHO chimaera in closed conformation, showing the N3 block helix (red) bound to β2 subunit (green).

**C** HeLa 1E cell lines, deleted for FCHO1 and FCHO2 ((Umasankar et al., 2014) transfected with constructs encoding GFP N-terminally appended to portions of FCHO2 as indicated. Transfection with GFP FCHOFBAR+wt linker (residues 1-416) rescues the fewer, large irregular AP2-containing CCPs patch phenotype caused by FCHO deletion and described in (*15*) (second row) to an increased number of standard sized CCPs. This is similar to transfection with full length GFPwtFCHO (lowest row right hand side control panels). Constructs comprising the BAR domain of FCHO only (residues 1-275) (top row) or comprising the BAR+ linker (residues 1-416) but with the linker N3 block deleted and replaced by an unstructured polypeptide (GFPFCHOBAR+linker ΔN3) (third row) fail to rescue the CCP phenotype as does transfection with GFP alone (lowest row left hand control panels and (*15*). GFP constructs are in green, AP2-containing clathirn-coated structures shown in red and the nucleus indicated in blue. These data indicate that the presence of N3 block of the linker seen by crystallography (Figs.3A and S4B) bound to β2 trunk is important for FCHO2 function in respect to AP2 binding and CCP morphology in vivo despite N3 being only one of four FCHO linker:AP2 contact sites and that on its own N3 displays only minimal binding so reaffirming the importance of avidity in the interaction: These data demonstrate transfer of data from structure back to cell biology

**D** Coomassie stained SDS PAGE gel of AP2:µ2FCHO2linker following placing on grids. The upper band (~70kDa) of the chimaeric µ2 band shows no degradation and there is no 50kDa band corresponding to apo µ2 (i.e. with no appended FCHO linker): Together these imply there has been no degradation or cleavage of the construct in solution suggesting that the apparent absence of Cµ2 in ~17% of particles is due to its static disorder following its displacement from AP2 bowls once they have been transitioned into an open conformation due to FCHO linker induced destabilization

**E** Cryo-EM reconstruction (3.8 Å) of AP2:µ2FCHO2-N1+N2+N3 chimera majority class. The map is a result of a reconstruction of the highest uniform population of particles (Cµ2-in) (see also Fig. S5). The AP2 is in the closed conformation and is coloured according to the underlying subunits with the position of the FCHO linker N1 block indicated in yellow.

**F** Ribbon representation in same orientation and colour scheme as **E**. For orientation purposes only, N3, which is not seen in SPA is shown in red, in thin worm representation in the position defined by X-ray crystallography (Fig 3).

**G** Structure as in **F** overlayed with semi-transparent, white Cryo-EM reconstruction (3.8 Å).

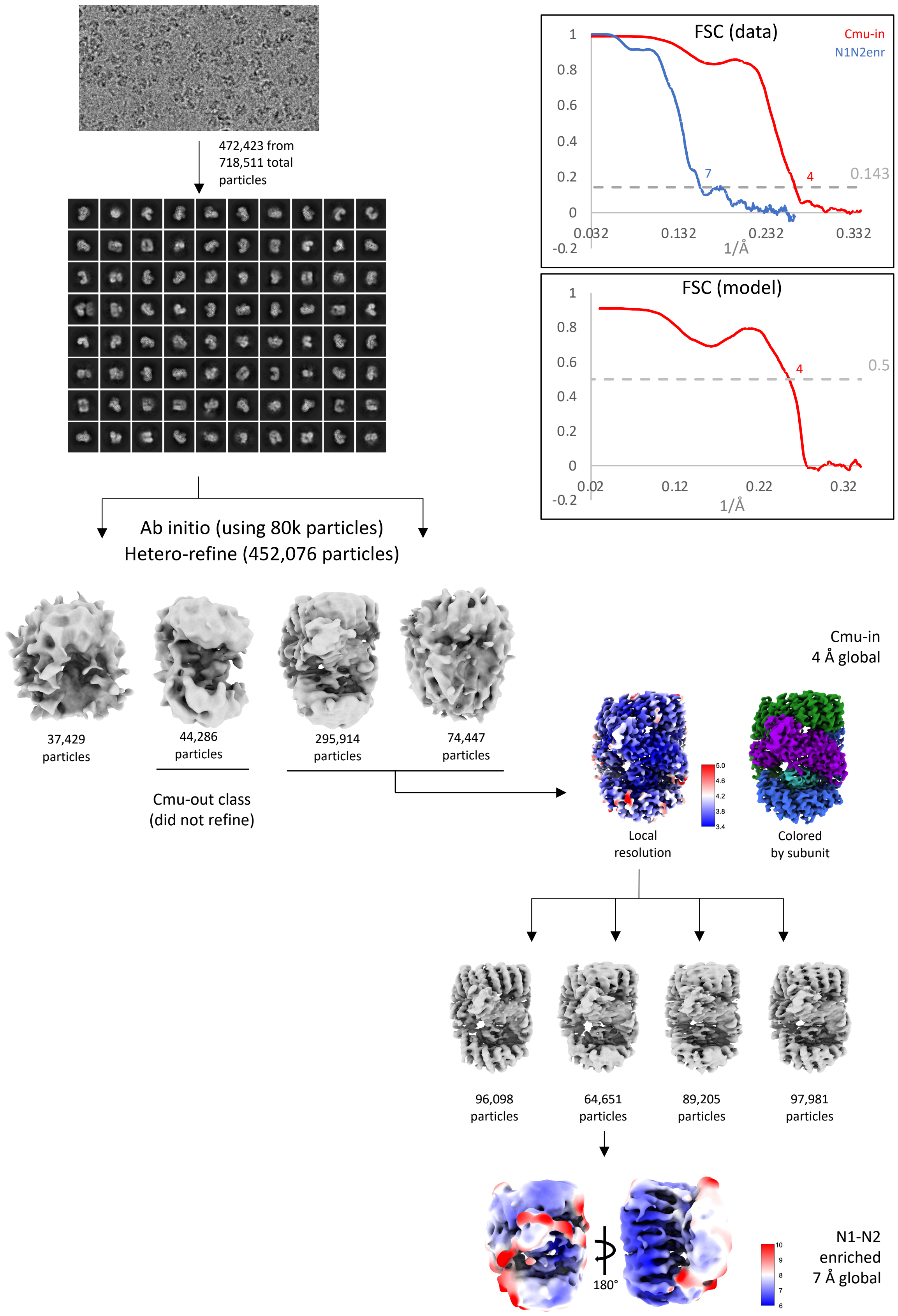

472,423 from
718,511 total
particles
FSC (data)
Cmu-in
N1N2enr
1
0.8
0.6
0.4
0.2
0
-0.2
7
4
0.143
0.032
0.132
0.232
0.332
1/Å
FSC (model)
1
0.8
0.6
0.4
0.2
0
-0.2
4
0.5
0.02
0.12
0.22
0.32
1/Å
Ab initio (using 80k particles)
Hetero-refine (452,076 particles)
37,429
particles
44,286
particles
295,914
particles
74,447
particles
Cmu-out class
(did not refine)
Cmu-in
4 Å global
5.0
4.6
4.2
3.8
3.4
Local
resolution
Colored
by subunit
96,098
particles
64,651
particles
89,205
particles
97,981
particles
180°
10
9
8
7
6
N1-N2
enriched
7 Å global

**Supplemental Figure S5. Single particle cryo-EM image processing workflow.**

AP2 particles were automatically picked and extracted from micrographs following motion and contrast transfer function correction. Particles were first subjected to 2D classification and classes with poor resolution or appearance were removed before *ab initio* models were determined using cryoSPARC and all particles were sorted into 4 classes. The majority class, a pool of two subclasses resembling closed AP2 core, was refined and also further sub-classified into 4 populations, each with small amounts of density outside of that expected for AP2 core alone both in the region of FCHO2 N1 and in the region adjacent on Cμ2. One class had density continuous with FCHO2 N1 (location determined by crystallography), shown here as the N1N2-enriched subclass. Further processing of both the overall Cμ2-in structure and the N1N2-enriched subclass involved particle reduction as mitigation for preferential orientation (see M&M) followed by non-uniform refinement. Reconstructions of the overall Cμ2-in structure and the N1N2-enriched structure are illustrated with estimated local resolution. Local filtering was applied to the refined Cμ2-out map after local resolution estimation. FSC curves for half-maps of each refined structure are shown in the top right panel, as well as that of the model vs data for the overall Cμ2-in structure.

Zaccai, Kadlecova Fig.S6

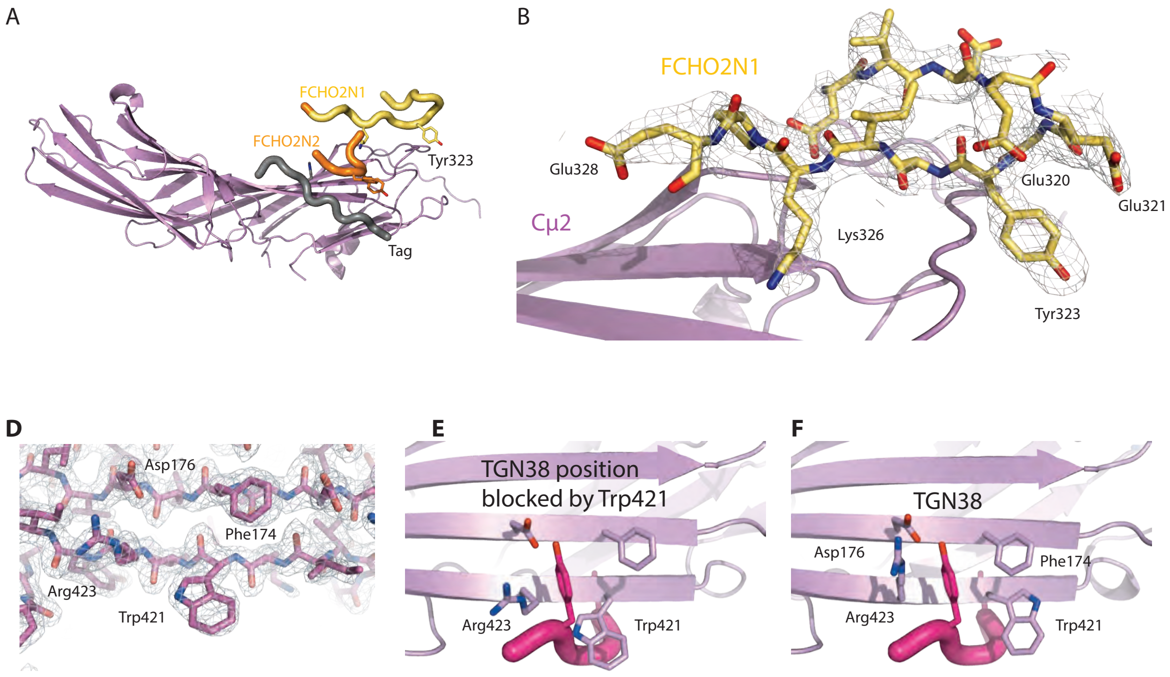

**Fig.S6 Structures of Cμ2:FCHO2-N1+N2 chimaeras**

**A** 2.3Å resolution overall structure of $His_6$-tagged Cμ2:FCHO2-N1+N2 chimaera: N1 (yellow) stays in the same position as in GST cleaved tag structure shown in Fig.4E: the main difference to cleaved GST tagged version is the position of the part of N2 (orange).

**B** 2.6 Å-resolution 2Fo-Fc electron density of Cμ2:FCHO2-N1+N2 chimaera (cleaved GST tag) contoured at 1.3σ showing molecular details of the binding of N1.

**D, E, F.** In the absence of YxxΦ ligand or a bound back affinity tag assuming its position, μ2Trp421 'swings back' into the hydrophobic Y pocket to shield it from solvent. **D** 1.9Å resolution 2Fo-Fc electron density contoured at 1.5σ showing μ2Trp421 'swung back' **E** compares this new position for μ2Trp421 with that of a standard YXXØ motif from published structures of complexes between Cμ2 and YXXØ motif peptides **F** : the μ2-Trp421 side chain is rotated through angles χ1 by ~110° and χ2 by ~130°, i.e. it is flipped back as compared to its position in a YXXØ motif liganded structures.

Zaccai, Kadlecova Fig.S7

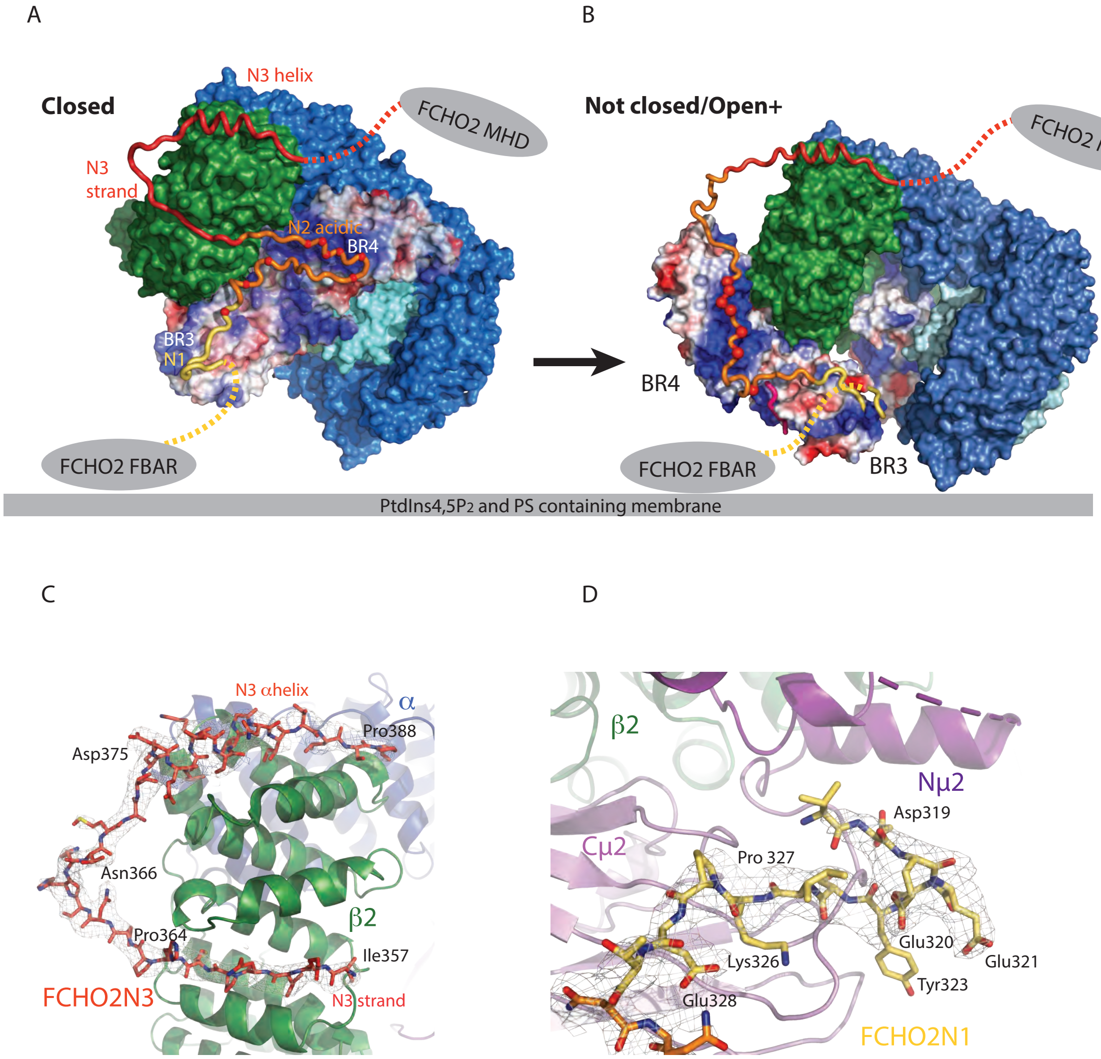

**Fig.S7 FCHO linker binds to β2 and μ2 subunits of AP2 using blocks N3 and N1 respectively**

**A** Model of the closed of AP2 core bound to FCHO2-N1N2N3. The FCHO2 blocks and the AP2 α, β2 and σ subunits are coloured as previously (N1 yellow, N2 orange, N3 red, α blue, β2 green and σ2 cyan). The Cμ2 is shown in electrostatic surface representation to demonstrate how the FCHO2 acidic region (acidic residues as red spheres) could interact with the positively charged surface of Cμ2. The closed AP2 is loosely bound to the PM only via it's a PtdIns4,5$P_2$-binding site.

**B** Once bound to closed AP2, the N1-N3 linker could destabilise the closed conformation by interfering with the Cμ2 / β2 subunit interface in the region where YxxΦ cargo binds to β2Val365 and β2Tyr405. Once the AP2 'bowl' changes shape the Cμ2 will be ejected and could take up a number of positions, which we believe are averaged out in our SPA (Fig. 3C) in which the YxxΦ cargo binding site is free (in line with data in Fig. S5) – one such possible conformation of AP2 is that which we have previously designated as Open+, which is modelled bound to FCHO2 linker here such that N, N2 and N3 can all bond simultaneously but Cμ2 is not membrane bound but AP2 is loosely bound to the PM via it's a PtdIns4,5$P_2$-binding site.

**C, D** 3.3Å resolution 2Fo-Fc electron density for the FCHO2 linker bound to open AP2 contoured at 1σ. (**C**) N3 block (red) bound to β2 subunit (green) (**D**) N1 (yellow) bound to Cμ2 (purple): important side chains in the FCHO linker side of the binding are indicated.

Zaccai, Kadlecova Figure S8

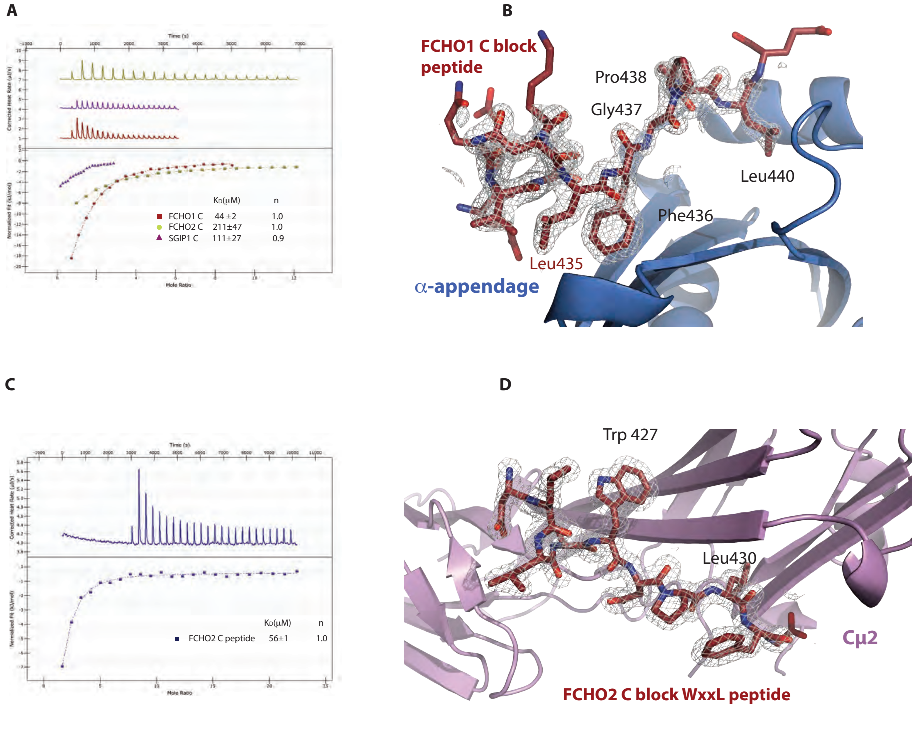

**Fig.S8 Details of FCHO C block binding to α-appendage and to Cμ2**

**A, B** FCHO1 C block (claret) bound to AP2 α-appendage (blue). **A** Isothermal Calorimetry analysis and resultant $K_D$s of C block peptides of FCHO1 (red $K_D$ ~40μM), FCHO2 (green $K_D$ ~200μM) and SGIP (purple $K_D$ ~110μM) binding to α-appendage. **B** 1.4 Å resolution 2Fo-Fc map contoured at 1.6σ. An apo structure of the α-appendage was used as MR search model (PDB 1W80 without ligands) – the direction of the peptide chain is clearly opposite to that of the published DP[FW], FxDxF motif/α-appendage complexes.

**C, D** FCHO2 C block (claret) in the YxxΦ binding site on Cμ2 (purple). **C** Isothermal Calorimetry analysis of FCHO2 C block peptide binding to AP2 Cμ2 Resultant $K_D$ ~50μM (blue): the tightest binding cargo YxxΦ peptide is TGN38 ($K_D$~2μM) with standard cargo binding having $K_D$s between 30 and 100μM (*54*). No binding was observed for FCHO1 and SGIP C block peptides. **D** 1.7Å resolution 2Fo-Fc map contoured at 1.6σ. An apo structure of Cμ2 was used as MR search model (PDB 1BXX without ligands).

Zaccai, Kadlecova Fig.S9

A

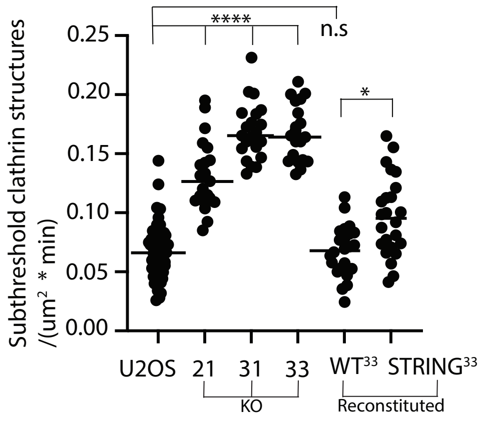

B

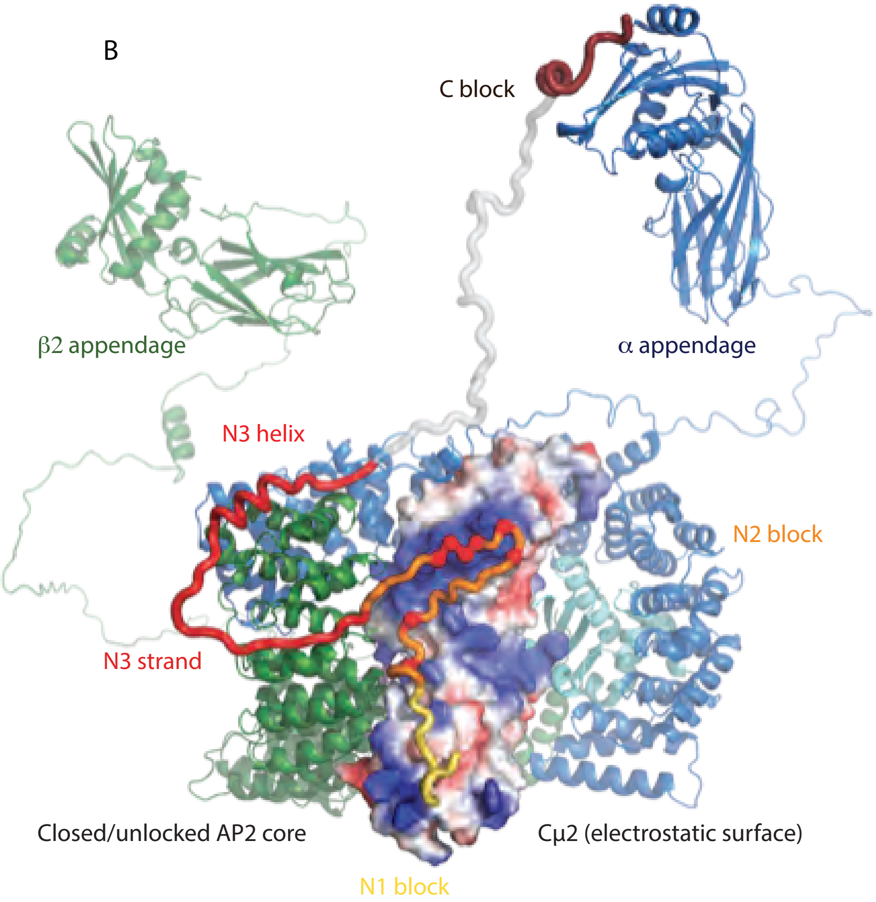

**Fig.S9 FCHO linker binding to full length AP2:cellular effect and molecular model**

**A** Comparison of initiation densities of transient dim clathrin coated structures for parental cell line, isogenic knockout clones 21,31 and 33, $FCHO2^{33\text{-}WT}$ and $FCHO2^{33\text{–}STRING}$ cell lines.

**B** Composite structural model of the full length AP2 in complex with FCHO2. The FCHO2 blocks and the AP2 $\alpha$, $\beta$2 and $\sigma$ subunits are coloured as previously (N1 yellow, N2 orange, N3 red, C block claret, $\alpha$ blue, $\beta$2 green and $\sigma$2 cyan). The C$\mu$2 is shown in electrostatic surface representation to demonstrate how the FCHO2 acidic region (acidic residues as red spheres) could interact with the positively charged surface of C$\mu$2. The length of the linkers separating the various boxes would allow all to bind simultaneously i.e. N1 and N2 binding to C$\mu$2, N3 binding to $\beta$2 and C binding to $\alpha$-appendage

Zaccai, Kadlecova Figure S10

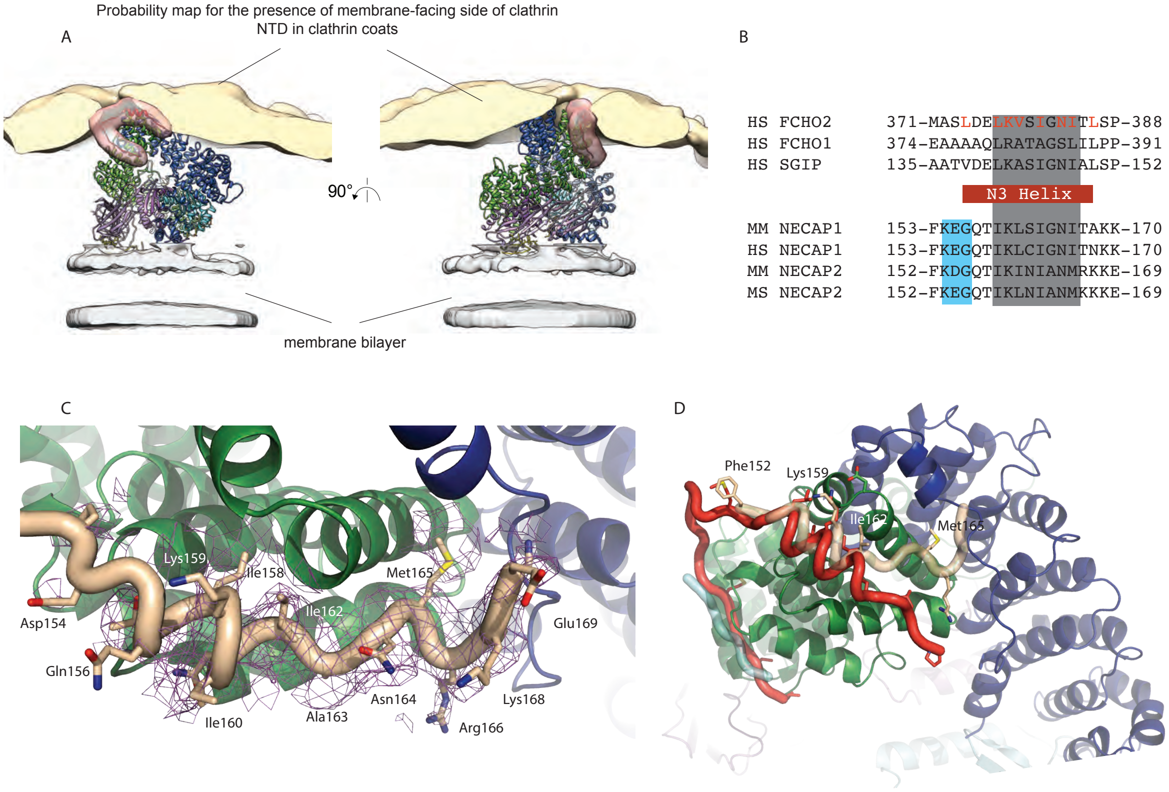

**Fig.S10 Competing interactions for FCHO linker blocks on AP2**

**A** The position of the FCHO2 linker relative to AP2 and Clathrin when assembled into a clathrin coat. Membrane-recruited AP2 is depicted as a ribbon model (PDB: 6YAF) and the corresponding EM density is cropped to only illustrate the bilayer region for clarity. The position of the FCHO2 linker was transferred from the X-ray structure of AP2/FCHO2 complex described in this study and is shown as a ribbon model in red and is overplayed with semi-transparent simulated density map derived from cryo EM tomography (*9*). The yellow volume illustrates the probability map of positions of the center of the membrane-proximal face of the clathrin NTDs relative to AP2, at an arbitrarily selected threshold corresponding to approximately 10% of relative subtomogram positions. The FCHO2 linker would be positioned adjacent to and partially overlapping with NTDs of the polymerized clathrin layer and so would be competed off AP2 by clathrin.

**B, C, D** Possible alternative model interpretation from (*57*) of NECAP2 “Ex” segment in complex with AP2 $\beta$2 subunit.

**B** Sequence homology between N3 helix (376-ELKVSIGNIT-385), mouse NECAP1 “Ex” segments (158-TIKLSIGNIT-167) and mouse NECAP2 “Ex” segment (157-TIKINIANMR-166) allowed NECAP2 “Ex” segment to be positioned in the N3 binding site of the AP2 $\beta 2$ subunit. Homology is indicated in grey, helical prediction in red and the conserved AP2 binding NECAP1 KEG motif is highlighted in blue (*57*)

**C** Structural comparison between FCHO2 N3 (red) and the alternative NECAP2 “Ex” (yellow) binding position to AP2 $\beta$2 (green) and $\alpha$ (blue) subunits built into the clearly helical density shown in C and refined locally using Coot. The key side chains from NECAP, which are conserved in property to the homologous stretch on N3 (shown in B), are indicated

**D** In the 3.5Å SPA cryo-EM structure of AP2 in complex with NECAP2 (purple mesh) (*57*), there is $\alpha$-helical density. The structure of NECAP2 can be readily built into this, on the basis of the FCHO NECAP sequence alignment in B and the density for the NECAP2 specific side chains of Met165 and Arg166 visible. The strongly conserved KEG motif can then be positioned to interact with $\beta$2, with the lysine's amine tentatively hydrogen bonding to the main chain carbonyl of $\beta$2-Lys494. Importantly, the double mutant KEG to AES abolishes NECAP Ex binding to AP2 (*63*)

If mammalian NECAP 'Ex' segment does indeed occupy the same position as FCHO N3 PAP2•NECAP complex, this would then allow the intervening NECAP main chain to run down the side of the $\beta$2 solenoid and would necessitate a repositioning of the KEG sequence originally defined in (*57*) as important for AP2 binding. In such a scenario, FCHO N3 and NECAP Ex would compete for the same binding site on $\beta$2 and this will impact their temporal ordering during CCV formation and help to explain their antagonistic actions (*57*). The new positioning of NECAP Ex helix may also have implications for the mechanistic model presented for NECAP function with regards to binding phosphorylated AP2 conformers (*48, 57*). NECAP's integration into CCP will likely be most efficient at the neck/edge of a Ω structure since it will also be connected to SNX9/Amphiphysins BAR domain-containing proteins, which have a preference for binding 'tubular' membrane structures of this curvature (*41*). However, as its Ex domain, like FCHO's N3, sterically overlaps with the clathrin TD layer, NECAP's binding to PAP2 could encourage PAP2 dissociation from the CCP by driving it from the clathrin lattice as well as probably shifting its equilibrium towards a closed non-membrane-attached form. The net result would be to exclude PAP2 and any attached cargo from the CCP's neck so both removing competition for PtdIns4,5$P_2$

for SNX9/Amphiphysin-recruited dynamin to bind and to allow complete neck construction, both of which would favour vesicle scission.

**Table S1. Cryo-electron tomography data collection.**

| | AP2 | AP2+FCHO |
|---|---|---|
| **Microscope, Voltage (keV)** | Titan Krios, 300 | |
| **Detector** | Gatan Quantum K3 | |
| **Energy filter slit width (eV)** | 20 | |
| **Electron exposure (e/Å$^2$) dose fractionation** | ~130, uniformly distributed over tilt series | |
| **Defocus range (μm)** | 1.0 – 3.5 | |
| **Tilt scheme (min/max, step)** | -60°/+60, 3°, dose-symmetrical (Hagen scheme) | |
| **Movie recording** | 10 frames per tilt | |
| **Magnification (times)** | X53,000 | |
| **Pixel size (Å)** | 1.701 | |
| **Number of tomograms acquired/used (no.)** | 18/17 | 32/22 |
| **Traced liposomes (no.)** | 140 | 238 |

**Table S2. Subtomogram averaging image processing parameters and statistics.**

| | AP2 | AP2+FCHO |
|---|---|---|
| **EMDB** | EMD-???? | EMD-???? |
| **EMPIAR** | EMPIAR-?????? | EMPIAR-?????? |
| **Particles count (no.):** | | |
| **Initial geometric seeding** | 160,263 | 290,187 |
| **After cross-correlation cleaning in bin8** | 112,945 | 269,280 |
| **After distance cleaning in bin4** | 44,514 | 95,039 |
| **After cross-correlation cleaning in bin4** | 41,215 | 86,311 |
| **After removal of empty classes** | 37,600 | 71,134 |
| **After cross-correlation cleaning in bin2** | 27,437 | 51,868 |
| **EM maps** | | |
| **Symmetry imposed** | none | |
| **Corrected resolution at FSC threshold 0.143 (Å)** | 11.7 | 9.5 |

**Table S3 Crystallographic data collection and refinement statistics**

| | AP2 (with β2-FCHO2 chimera) | AP2 in complex with FCHO2linker and TGN | AP2 in complex with SeMetFCHO2 and TGN |
|---|---|---|---|
| **PDB** | 7OHO | 7OG1 | - |
| **Space group** | P $3_1$ 2 1 | P $2_1$ | P $2_1$ |
| **Cell dimensions (Å, °)** (a, b, c, $\alpha$, $\beta$, $\gamma$) | 122.0, 122.0, 257.4, 90, 90, 120 | 92.6, 150.0, 96.4, 90, 112.7, 90 | 92.0, 146.8, 99.5, 90, 116.4, 90 |
| **Resolution range (Å)** (outer shell) | 66.61-2.88 (2.95-2.88) | 78.58-3.25 (3.39-3.25) | 76.08-3.97 (4.35-3.97) |
| **$R_{merge}$** | 0.115 (2.019) | 0.133 (2.123) | 1.109 (4.455) |
| **$R_{pim}$** | 0.037 (0.649) | 0.061 (1.049) | 0.458 (1.840) |
| **Number of observations** | 500,376 | 253,506 | 276,139 |
| **Number of reflections** (outer shell) | 51,118 (3,710) | 38,355 (4,690) | 20,527 (4,887) |
| **<(I)/$\sigma$(I)>** | 13.5 (1.2) | 7.2 (0.8) | 2.7 (1.0) |
| **CC(½)** | 0.998 (0.529) | 0.998 (0.366) | 0.956 (0.502) |
| **Completeness (%)** | 99.9 (99.9) | 100 (100) | 100 (100) |
| **Multiplicity** | 9.8 (9.6) | 6.6 (6.0) | 6.8 (6.8) (anomalous) |
| **Wilson B factor (Å$^2$)** | 80 | 102 | 62 |
| **Resolution (Å)** | 61.02 - 2.88 | 76.54 - 3.25 | - |
| **Reflections** (total / test) | 51,118 / 2,547 | 38,355 / 1,881 | |
| **R / $R_{free}$ (%)** | 18.6 / 26.9 | 20.8 / 31.4 | - |
| **Average B factor (Å$^2$)** (all / AP2 / FCHO / other) | 104 / 91 / 176 / 110 | 145 / 144 / 166 / 177 | - |
| **RMS bond lengths (Å)** | 0.008 | 0.006 | - |
| **RMS bond angles (°)** | 1.606 | 1.497 | - |
| **Ramachandran (%)** (favored / outliers) | 87 / 3 | 90 / 2 | |

**Table S4 Crystallographic data collection and refinement statistics**

| | **$His_6$-Cμ2 (apo)** | **$His_6$-Cμ2-FCHO2** (His6-tagged) | **Cμ2-FCHO2** (cleaved GST) |
|---|---|---|---|
| **PDB** | 7OFP | 7OHZ | 7OI5 |
| **Space group** | $P\ 6_5$ | $P\ 2_1$ | C 2 |
| **Cell dimensions (Å, °)** (a, b, c, α, β, γ) | 123.8, 123.8, 112.7, 90, 90, 90 | 55.7, 129.9, 64.4, 90, 102.5, 90 | 116.0, 55.4, 167.7, 90, 109.0, 90 |
| **Resolution range (Å)** (outer shell) | 77.68–1.92 (1.95-1.92) | 62.90-2.27 (2.31-2.27) | 79.27-2.61 (2.65-2.61) |
| **Number of observations** | 1,321,253 | 253,807 | 203,739 |
| **Unique reflections (non-anomalous)** | 74,711 (3,642) | 40,099 (1,520) | 31,089 (1,543) |
| **Completeness (%)** | 99.9 (98.0) | 97.1 (72.4) | 99.9 (99.3) |
| **Multiplicity** | 17.7 (12.1) | 6.3 (4.5) | 6.6 (6.2) |
| **$\langle I/\sigma(I) \rangle$** | 13.1 (1.0) | 7.0 (1.2) | 6.6 (1.0) |
| **$R_{merge}$** | 0.223 (6.379) | 0.136 (1.165) | 0.163 (1.576) |
| **$R_{pim}$** | 0.055 (1.912) | 0.058 (0.593) | 0.068 (0.693) |
| **CC(½)** | 0.999 (0.440) | 0.996 (0.602) | 0.995 (0.374) |
| **Wilson B factor ($Å^2$)** | 31 | 38 | 52 |
| **Resolution (Å)** | 77.68 - 1.92 | 62.90 - 2.27 | 79.27 - 2.61 |
| **Reflections** (total / test) | 74,663 / 3,819 | 38,073 / 1,989 | 30,964 / 1,557 |
| **$R$ / $R_{free}$ (%)** | 18.5 / 20.7 | 27.0 / 31.7 | 24.5 / 29.4 |
| **Average B factor ($Å^2$)** (all / AP2 / FCHO / other) | 42 / 42 / - / 45 | 54 / 54 / 65 / 44 | 73 / 72 / 93 / 53 |
| **RMS bond lengths (Å)** | 0.012 | 0.003 | 0.003 |
| **RMS bond angles (°)** | 1.060 | 0.643 | 0.683 |
| **Ramachandran (%)** (favored / outliers) | 96 / 1 | 94 / 0 | 94 / 0 |

**Table S5 Crystallographic data collection and refinement statistics**

| | α ear with FCHO1 C block | $His_6$-Cμ2 with FCHO2 C block | $His_6$-Cμ2 with FCHO2 C block |
|---|---|---|---|
| **PDB** | 7OHI | 7OIT | 7OIQ |
| **Space group** | C 2 2 $2_1$ | P $3_2$ 2 1 | C 2 |
| **Cell dimensions (Å, °)** (a,b,c, α,β,γ) | 61.2, 145.5, 88.8, 90, 90, 90 | 66.4, 66.4, 161.6, 90, 90, 120 | 118.9, 64.5, 108.4, 90, 112.0, 90 |
| **Resolution range (Å)** (outer shell) | 47.60-1.41 (1.43-1.41) | 57.47–1.65 (1.68–1.65) | 55.70–1.85 (1.88–1.85) |
| **Number of observations** | 873,264 | 968,797 | 354,995 |
| **Unique reflections** (outer shell) | 73,320 (2,558) | 50,587 (2,463) | 58,302 (1,599) |
| **Completeness (%)** | 95.9 (67.9) | 100.0 (100.0) | 89.4 (49.1) |
| **Multiplicity** | 11.9 (6.4) | 19.2 (17.0) | 6.1 (4.0) |
| **<(I)/σ(I)>** | 17.0 (0.2) | 19.0 (1.1) | 9.7 (1.1) |
| **$R_{merge}$** | 0.059 (2.810) | 0.082 (2.696) | 0.091 (1.064) |
| **$R_{pim}$** | 0.017 (1.157) | 0.019 (0.668) | 0.039 (0.577) |
| **CC(½)** | 1.000 (0.253) | 1.000 (0.462) | 0.995 (0.476) |
| **Wilson B factor (Å$^2$)** | 23 | 27 | 30 |
| **Resolution (Å)** | 47.60 - 1.41 | 57.47 - 1.65 | 55.70 -1.85 |
| **Reflections** (total / test) | 71,365 / 3,608 | 47,975 / 2,538 | 58,280 / 2,932 |
| **R / $R_{free}$ (%)** | 20.3 / 21.9 | 19.5 / 21.5 | 19.5 / 21.7 |
| **Average B factor (Å$^2$)** (all / AP2 / FCHO / other) | 33 / 32 / 55 / 43 | 35 / 34 / 31 / 44 | 40 / 40 / 42 / 45 |
| **RMS bond lengths (Å)** | 0.006 | 0.013 | 0.011 |
| **RMS bond angles (°)** | 0.905 | 1.801 | 1.658 |
| **Ramachandran (%)** (favored / outliers ) | 98 / 0 | 98 / 0 | 97 / 0 |

**Table S6. Cryo-electron microscopy data collection, refinement and validation statistics for single-particle structures of AP2:β2FCHO2linker chimaera in solution.**

| | | |
|---|---|---|
| Voltage (keV) | 300 | |
| Microscope and Detector | Titan Krios, K3 | |
| Energy filter slit width (eV) | 20 | |
| Electron exposure (e/ Å2) | 47.28 | |
| Defocus range (μm) | 0.8-2.8 | |
| Movie recording | 48 frames | |
| Magnification | 130,000 | |
| Pixel size (Å) | 0.326 | |
| Initial number of particles (number used for refinement) | 718,511 (472,423) | |
| | **CMu-in** | **CMu-in N1+N2** |
| EMDB/PDB ID | XXXX/XXXX | XXXX/XXXX |
| Final number of particles (no.) | 143,815 | 28,392 |
| FSC data (0.143) (Å) | 3.8 | 6.7 |
| FSC model (0.500) (Å) | 3.9 | |
| Initial model, PDB ID | 2vgl | 2vgl + N1N2 |
| Non-hydrogen atoms | 13,707 | 13,982 |
| Protein residues | 1717 | 1751 |
| Ligands | 0 | |
| R.m.s. deviations | | |
| Bond length (Å) | 0.003 | 0.004 |
| Bond angle (°) | 0.549 | 0.820 |
| Validation | | |
| Molprobity score | 1.74 | 2.42 |
| Clashscore | 9.53 | 28.75 |
| Poor rotamers (%) | 0 | 0 |
| Ramachandran plot | | |
| Favoured (%) | 96.42 | 92.33 |
| Allowed (%) | 3.58 | 7.61 |
| Outliers (%) | 0.0 | 0.06 |

**Supplementary Videos**

**Video S1** Super-resolution live-cell imaging of CME dynamics by eTIRF-SIM in U-2 OS cells expressing m-Scarlet-FCHO2$^{WT}$ and eGFP-CLCa. Movies were acquired in sequential mode for 6 minutes with 2s frame intervals.

**Upper panel:** Averaged TIRF time-lapse movie: For each channel the nine frames of raw TIRF–SIM data were gathered with three different phases and angles at every time point and then averaged for CCP detection and tracking with CMEanalysis analysis suite and custom Matlab scripts. The resulting detected spatial coordinates of valid tracks were superimposed on the final eTIRF-SIM reconstructed time-lapse movie (Lower panel).

**Video S2** Final eTIRF-SIM reconstructed time-lapse movie of eGFP-CLCa in U-2 OS cells.

**Video S3** Final eTIRF-SIM reconstructed time-lapse movie of mScarlet-FCHO2$^{WT}$

**Video S4** Formation of a single CCP imaged by eTIRF-SIM for 65 seconds with 2s frame interval**.** Initiation, growth, and scission of were visualized with eGFP-CLCa.

**Video S5**. Conformational variability in single particle EM reconstruction of AP2 FCHO2 chimera. 3D variability analysis (3DVA; CryoSPARC) was carried out on the majority dataset of AP2 FCHO2 chimera in which the stacked [illegible]-solenoid structures of AP2 contract with a concurrent elongation of the long axis of the AP2 core particle. This motion may be a step towards opening of the AP2 bowl, wherein Cμ2 is ejected and the α and β2 trunks converge by a distance of ~16Å.
